# Ecological divergence of sympatric *Saccharomyces* species across wild and fermentative environments in the neotropics

**DOI:** 10.1101/2025.05.31.656962

**Authors:** Claudio López-Gallegos, Xitlali Aguirre-Dugua, J. Abraham Avelar-Rivas, Manuel R. Kirchmayr, Lucía Morales, Alexander DeLuna, Eugenio Mancera

## Abstract

The common yeast *Saccharomyces cerevisiae* is widely associated with anthropogenic habitats and dominates agave fermentations in Mexico, despite the absence of starter cultures in artisanal production. Yet, the origins and dispersal of the microorganisms involved in this fermentation system—including wild *S. cerevisiae* and its sister species *S. paradoxus*—remain poorly understood. Here, we analyzed the distribution of these two species across three regions in Mexico by collecting a total of 861 samples from 15 traditional distilleries and 25 nearby natural sites, including tree bark, insects, fermentation tanks, and distillery surfaces. Among 4,006 isolates identified by MALDI-TOF biotyping, *S. cerevisiae* and *S. paradoxus* were recovered at overall rates of 21% and 3.4%, respectively, with the former more frequently found in distilleries and the latter in natural habitats. Notably, *Saccharomyces* isolates were not exclusively associated with oak trees, as strains were also recovered from leguminous, pine, and Anacardiaceae trees, suggesting that the oak specialization observed in temperate regions may not hold in tropical environments. Genome sequencing of 100 *Saccharomyces* isolates and SNP-based analyses showed that *S. cerevisiae* strains clustered by geography, not by substrate, and were largely confined to two Mexican Agave clades, with minor representation from the North American Oak, Pacific West Coast, and Wine clades. Strains isolated from agave fermentations belonged to the same populations as those from plant substrates, with insects, especially Drosophilidae, likely mediating their dispersal. Meanwhile, *S. paradoxus* isolates grouped into the SpA (Eurasian) and SpB (North American) lineages, with three novel SpB subclades identified. Intriguingly, one of these, the SpB_MxAgave population, was exclusively associated with agave fermentations, providing the first evidence of a persistent association of *S. paradoxus* with an anthropogenic habitat. Our study sheds light on the ecological dynamics and diversity of *S. cerevisiae* and *S. paradoxus* populations in the neotropics.

## Introduction

*Saccharomyces* yeasts (Saccharomycotinae) are fundamental for alcoholic fermentation in traditional and industrial food production systems worldwide. Among them, the common yeast *S. cerevisiae* is mostly found in anthropogenic fermentative environments related to the production of bread, beer, wine, dairy products, sake, and many other foods and beverages of local importance (Alsammar & Delneri, 2020). In the last ten years, different studies have provided insight into the biogeography of *S. cerevisiae*, revealing population lineages that correlate with world regions and the environmental and fermentation habitats of the species. These findings are starting to reveal the evolutionary relationships among domesticated and wild strains (Han *et al*., 2021; Loegler, 2024; Peter *et al*., 2018; Pontes *et al*., 2020).

Wild populations of *S. cerevisiae* were formerly thought to be rare or nonexistent, but more extensive sampling and an increasing interest in yeast ecology in natural ecosystems have shown that wild *S. cerevisiae* can be found in temperate forest habitats, specifically on soil, decayed wood and tree bark (Almeida *et al*., 2015; Bai *et al*., 2022; Barbosa *et al*., 2016; Spurley *et al*., 2021). According to phylogenetic analyses, some wild strains are distantly related to and have remained isolated from human-associated populations (e.g., wild strains from Taiwan and China; Loegler 2024). In other cases, wild strains show sister-relationships with domesticated strains and, as a result, have been inferred to be the ancestral population from which domesticated stocks descend (for instance, Mediterranean Oak and Wine/European strains; Almeida *et al*., 2015). Following the complex evolutionary relationships observed among strains, different domestication trajectories have been inferred along the *S. cerevisiae* phylogeny (Peter *et al*., 2018; Pontes *et al*., 2020). The widespread occurrence of strains with mosaic genomes has also shown that genetic admixture plays an important role in the evolution of *S. cerevisiae* populations and their adaptation to different environments (Barbosa *et al*., 2016; Liti *et al*., 2009; Marr *et al*., 2023; Pontes *et al*., 2020).

In contrast to *S. cerevisiae*, its sister species *S. paradoxus* is known to principally inhabit tree bark and soil in natural environments, displaying a stronger geographic structure with greater divergence levels across continents (Boynton & Greig, 2014; Liti *et al*., 2009). Populations of *S. paradoxus* from Eurasia are grouped in the SpA lineage, whereas the North American populations include SpA, SpB and SpC lineages, as well as their hybrid derived SpC* and SpD clades (Eberlein *et al*., 2019). The SpB lineage has also been identified in South America (Borneman & Pretorius, 2015). The co-occurrence of *S. paradoxus* and *S. cerevisiae* in the same temperate arboreal habitats from the Mediterranean, Central Europe and North America has led to the hypotheses that oak tree bark (genus *Quercus*, family Fagaceae) is their natural primary reservoir (Sampaio & Gonçalves, 2008; Sniegowski *et al*., 2002) and that contrasting temperature tolerances between the two species is the main driver of ecological differentiation at a local scale (Dashko *et al*., 2016; Gonçalves *et al*., 2011). However, these hypotheses are based on observations mainly done in temperate areas and the biogeography and ecology of these species in tropical regions remain unclear.

The genomic diversity of *Saccharomyces* yeasts in the tropical Americas has only recently begun to be thoroughly investigated, particularly in populations from South America (Barbosa *et al*., 2016; Barbosa *et al*., 2018). In the past years, our team described the rich yeast community from agave fermentations in Mexico, which is characterized by the ubiquity of *S. cerevisiae* and, surprisingly, the presence of *S. paradoxus* at low frequency (Gallegos-Casillas *et al*., 2024). Initial phylogenetic analyses of a broad set of *S. cerevisiae* strains from this environment showed that most form a monophyletic clade that is sister to the French-Guiana lineage, confirming previous relationships with other world’s lineages (Avelar-Rivas *et al*., 2024).

The agave fermentation environment represents a compelling system for analyzing *Saccharomyces* ecology and evolution because traditional fermentation of cooked agave for spirits production depends on microorganisms reaching the tanks “spontaneously” from the surrounding environment, since producers don’t use microbial inoculants. Moreover, many distilleries are small to medium-sized family-run units with seasonal production, during which fermentations are completely halted for several months (Arellano-Plaza *et al*., 2022). However, the origins and means of transportation of the microorganisms that are responsible for agave fermentations remain largely unknown. André Lachance (Lachance, 1995) addressed this question through taxonomic studies of the yeasts in a single distillery, suggesting that objects used during spirits production could serve as reservoirs and that *Drosophila* flies may act as vectors. Indeed, in wine open fermentations from Europe, *S. cerevisiae* can use insects such as wasps as reservoir and vectors, enabling its transport from the surrounding vineyard environment to the fermentation vats (Stefanini *et al*., 2012). More broadly, wasps, honeybees, flies and fruit flies have all been shown to successfully transport yeasts (Meriggi *et al*., 2020).

Agave fermentation systems in Mexico offer an excellent case study for examining the differential distribution of *S. cerevisiae* and *S. paradoxus* across natural and anthropogenic environments in a tropical region of the world. This system provides a unique opportunity to identify local reservoirs and to test whether yeast populations from natural and human-associated environments are connected through insect-mediated dispersal. In this study, we isolated wild *S. cerevisiae* and *S. paradoxus* strains from trees and insects in natural areas surrounding agave-producing regions. Comparing these isolates with those from agave fermentations, we found that *S. cerevisiae* was far more frequent in anthropogenic environments, whereas *S. paradoxus* was more common in natural habitats. Genome-wide phylogenetic analyses revealed distinct population-level distribution patterns. The same *S. cerevisiae* populations were detected in fermentation tanks, plants, tools, and insects within distilleries, as well as in natural substrates from nearby environments. In contrast, *S. paradoxus* strains were primarily isolated from natural substrates, though they were also found in agave fermentations. Remarkably, one *S. paradoxus* subclade was exclusively associated with agave fermentations. Altogether, our study sheds light on the ecological structure and diversity of *S. cerevisiae* and *S. paradoxus* populations in tropical regions of the world.

## Results

To better understand the interplay between natural and human-associated populations of *S. cerevisiae* and *S. paradoxus* in a tropical, megadiverse region of the world, we focused on three of the regions producing traditional agave spirits in Mexico: Southcentral (Oaxaca state), West I (Durango state) and Northeast (Nuevo León and Tamaulipas states). Overall, the most distant sampled locations are 1,137 km apart and the elevation range covered from 346 to 2,857 meters above sea level (**Fig. 1**). It is worth mentioning that although the sampled sites in the Northeast region were relatively close to each other, sites in the state of Nuevo León were on the Sierra Madre Oriental mountain range while the ones in the state of Tamaulipas were at the east of the range below the Sierra Madre. This is important because this range is known to represent a natural barrier for many plant taxa distributed across these two states (**Fig. 1C**).

**Figure 1.**
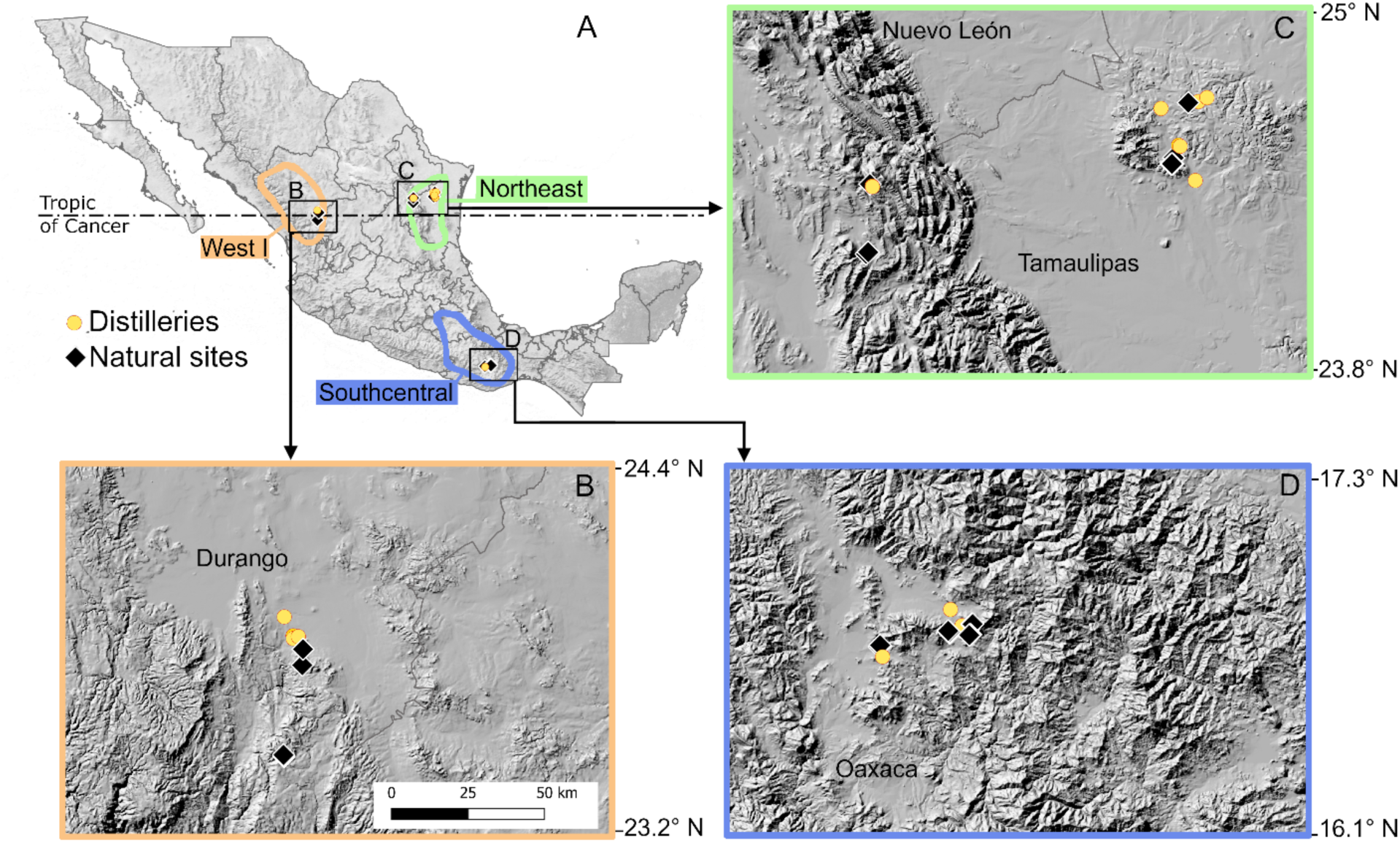
Sites sampled in three distinct agave spirit-producing regions of Mexico. **(A)** Overall location of the sites, including traditional distilleries (yellow circles) and sites in natural areas (black diamonds). The three producing regions are highlighted in color. Black squares indicate the areas shown in close-ups. Panels B to D show close-ups at equivalent scale for sampling areas in the West I **(B)**, Northeast **(C)**, and Southcentral **(D)** regions, respectively; gray lines indicate state boundaries.

In these three regions, we visited a total of 15 distilleries where samples were taken from tanks with active fermentations, bark from living trees and insects, as well as different objects used in the production process (tools, cooked agave stems, bottom and walls of empty vats, and fermentation waste, all of which we will refer to as “objects”). We also sampled 25 sites in natural areas near the distilleries, where bark from trees, insects and some fruits were collected (**Fig. 1**; **Suppl. Mat. S1**).

From the three regions we obtained a total of 861 samples from which we generated a collection of 4,006 yeast isolates by enriching with an isolation protocol designed to grow *Saccharomyces* yeasts (**Table 1**). From these isolates, 2,903 (73.4%) could be assigned a species with a MALDI-TOF Biotyper®, although only 2,209 (55.14%) were identified with confidence (score > 1.5). Using a stringent identification cutoff, 843 isolates (38.2%) were identified as *S. cerevisiae* and 136 (6.2%) as *S. paradoxus*, with no other *Saccharomyces* species detected. *Saccharomyces* isolates were recovered from all sampled distilleries and from 64% of the natural sites (16 out of 25).

**Table 1.** Number of samples per location type from different substrates and the resulting *Saccharomyces* spp. sequenced strains.

| Location Type | # Samples and *Saccharomyces* spp. isolation rate per substrate * |  |  |  |  | # Genome-sequenced strains |  |
| --- | --- | --- | --- | --- | --- | --- | --- |
|  | Total | Insects | Plants | Objects | Tanks | *S. cerevisiae* | *S. paradoxus* |
| Distilleries | 274 (41.87%) | 97 (27.83%) | 55 (18.19%) | 82 (47.56%) | 40 (85%) | 23 | 21 |
| Natural sites | 587 (5.45%) | 244 (2.05%) | 327 (7.34%) | - | - | 19 | 37 |
| Total | 861 | 341 | 382 | 82 | 40 | 42 | 58 |

\*Isolation rates are indicated in parentheses

Despite using an enrichment protocol specifically tailored for *Saccharomyces* species, a variety of non-*Saccharomyces* yeasts were isolated from both location types. The most frequently recovered species were *Lachancea thermotolerance*, *Pichia manshurica, Pichia kudriavzevii,* and *Kodamaea ohmeri* (**Suppl. Mat. S2**). Considering the different taxa identified in each location type, natural sites were more diverse (39 species, Shannon’s diversity index *H’* = 2.47) and less uniform (Simpson’s index *D* = 0.86) than distilleries (12 species, *H’* = 1.85, *D* = 0.67). Together, these results suggest that the agave fermentation system, considered within its natural context, offers a useful framework to investigate the ecological dynamics of *Saccharomyces* populations in the tropics.

### Differential distribution of *Saccharomyces* species between wild and anthropogenic environments

To assess possible differences in the distribution of the two *Saccharomyces* species that inhabit agave fermentations and the surrounding natural areas, we first estimated the success isolation rate as the percentage of the total number of samples from which *Saccharomyces* spp. isolates were recovered. The overall isolation rate of *Saccharomyces* spp. (i.e., considering both species) was 42% in distilleries and 5.5% in natural sites.

At the species level, *S. cerevisiae* was by far more abundant than *S. paradoxus* in distilleries (isolation rate 39.78% vs. 5.47%), while in natural areas they were both isolated at similar rates (3.92% vs. 3.74%, **Fig. 2A**). Compared to *S. cerevisiae, S. paradoxus* was a relatively rare species in distilleries, in agreement with our previous findings studying a larger set of agave fermentations (Gallegos-Casillas et al, 2024). A total of 746 *S. cerevisiae* strains were isolated from distilleries and 97 from natural sites, whereas 25 *S. paradoxus* strains were isolated from distilleries and 111 from natural sites. This difference in isolation frequencies between *S. cerevisiae* and *S. paradoxus* across natural and anthropogenic environments was statistically significant (Fisher’s Exact test, *p* = 2.2e–16), supporting a differential distribution of the two species by location type.

**Figure 2.**
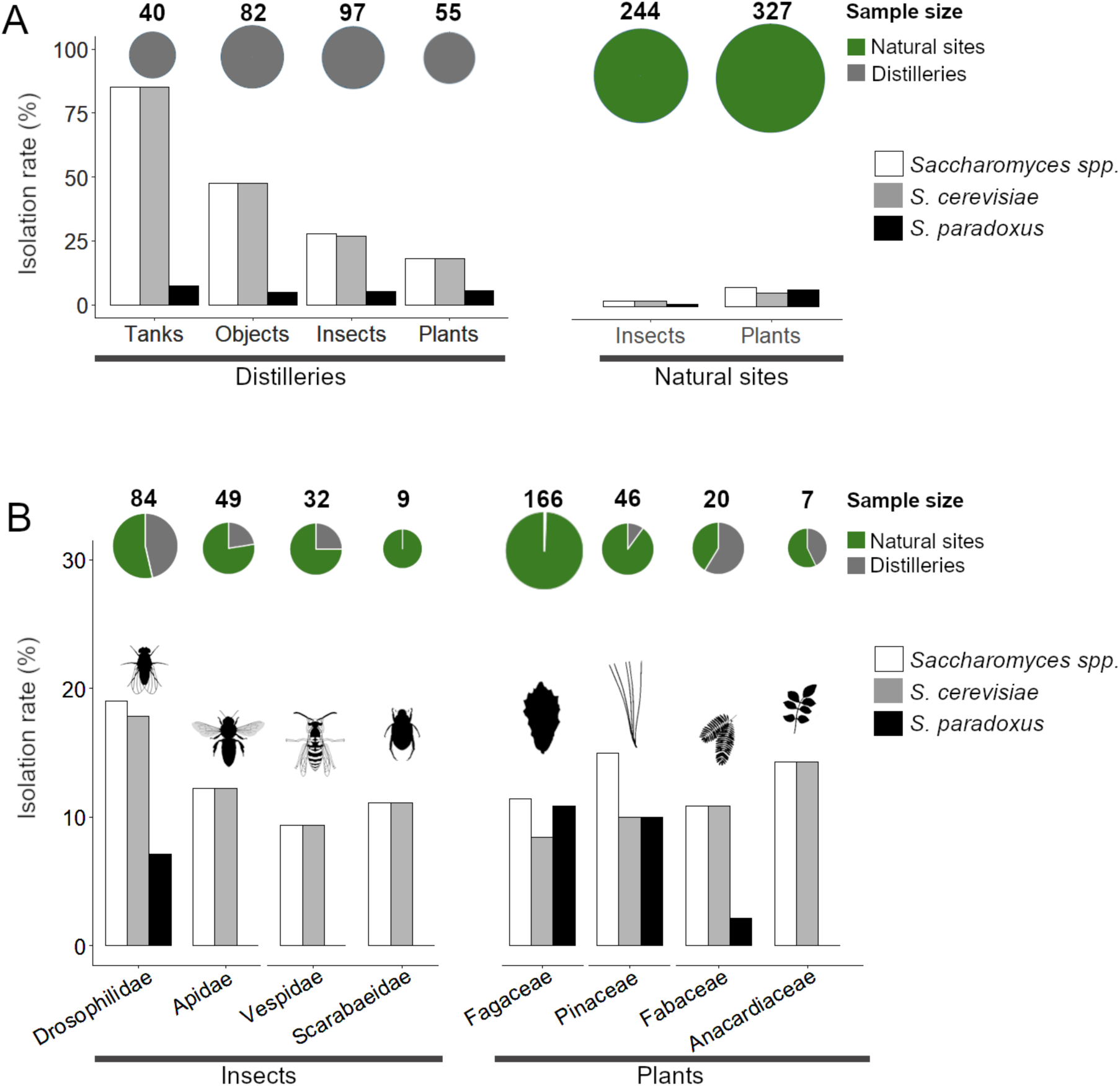
Isolation of *Saccharomyces* spp. from three geographical areas of Mexico. **(A)** Isolation rate of *Saccharomyces* spp. confidently identified with MALDI-TOF (score > 1.5) from each location and substrate type. **(B)** Isolation rate from different insect families (left) and plant families (right). Sample size of each category is shown on top. Isolation rate of *Saccharomyces* spp. (white bars) refers to the percentage of samples that yielded one or both species; samples yielding both species are counted only once.

In terms of substrates, within distilleries, we found that *S. cerevisiae* was more frequently isolated from tanks and objects, than from insects or plants. In contrast, *S. paradoxus* had a similar isolation rate among substrates within each location type and between distilleries and natural sites (**Fig. 2A**). In natural environments, most *Saccharomyces* isolates were isolated from plants, mostly from oak bark. We collected 165 bark samples from oaks in natural environments (and one from distillery), recovering 92 *S. paradoxus* and 59 *S. cerevisiae* isolates from 29 of them. Notably, twelve oak samples (7.3%) gave rise to both *Saccharomyces* species. However, irrespective of location type (distillery or natural site), isolation rate from oak trees was close to the average rate from other plant families (**Fig. 2B**; **Suppl. Table S3**). This was unexpected, given that oak trees have been reported as the preferred habitats for *Saccharomyces* species in temperate regions.

In the case of insects in natural environments, five samples of fruit flies (Drosophilidae) also hosted both species. Fruit flies were the insects from which more *Saccharomyces* isolates were recovered, and we also retrieved *S. cerevisiae* strains from wasps, bees and beetles, although no *S. paradoxus* isolates were retrieved from these other insect families (**Fig. 2B**). In summary, our results show that although both species are present across all sample types and occasionally co-occur, *S. cerevisiae* is substantially more common in distilleries, whereas *S. paradoxus* predominates in natural environments.

### *S. cerevisiae* populations are shared between distilleries and natural sites within regions

Our findings showed that natural environments surrounding distilleries harbor the same *Saccharomyces* species found in traditional agave fermentations. However, it remains possible that distinct populations of the same species inhabit each environment. To uncover the phylogenetic relationship of the *S. cerevisiae* strains isolated in different locations and substrates, we sequenced the genomes of 42 representative isolates (**Table 1**). These included 19 strains from natural sites and 23 from distilleries, selected based on the accuracy of MALDI-TOF identification, and the location type and geographical region of isolation. All *S. cerevisiae* sequenced strains from natural environments came from seven samples: three insects, three oaks and one leguminous tree (**Suppl. Table S1**). As detailed in Material and Methods, sequencing reads were aligned to a reference genome to identify SNPs in each strain. For phylogenetic analyses, we included genomic information from 56 strains covering the *S. cerevisiae* diversity from agave fermentations from the same three geographical areas and the state of Jalisco (West II region) (Avelar-Rivas *et al*., 2024), and 147 strains representative of the main world clades including strains from the originally named “Mexican Agave” clade (**Suppl. Table S1**). This resulted in a final matrix of 203 total strains and 870,190 SNPs from which we were able to clearly reconstruct the two main clades associated to traditional fermentations of agave from Mexico (Mexican Agave 1 and Mexican Agave 2), as well as their relationships with other closely related clades within the Neotropical cluster (Avelar-Rivas *et al*., 2024).

Overall, the phylogenetic tree of *S. cerevisiae* showed no associations between the phylogenetic placement of the strains and the substrate (insect, plant, object or tank) from which they were isolated (**Fig. 3A**). For example, isolates from insects clustered with strains from plants, from objects used during the production process and from the fermentations themselves, provided that they all were from the same geographical region. In terms of the location type, the nine strains from natural sites were grouped with strains from their same geographical region, isolated from fermentation tanks and other substrates within distilleries. For example, two isolates that were collected in the Southcentral region from an oak tree and a Diptera insect, grouped with strains isolated from insects and active fermentations from a distillery that is approximately ten km apart. The same occurred in the West I region, where two strains isolated from a beetle were closely related to a strain from a cooked agave stem sampled within a distillery at a similar distance range. These observations support the idea that plants and insects outside distilleries and far from fermentation activities host the same populations of *S. cerevisiae* associated with traditional open agave fermentations. This pattern was particularly evident for strains belonging to the Mexican Agave 2 and North American Oak clades. However, despite our sampling efforts, we were not able to isolate strains that belong to the Mexican Agave 1 clade from natural environments. All the strains in this clade were isolated in distilleries in the Northeast region and specifically in the state of Tamaulipas. Natural isolates from this producing region belong to the North American Oak clade and all were isolated in the state of Nuevo León on the Sierra Madre Oriental mountain range (**Fig. 1C**).

**Figure 3.**
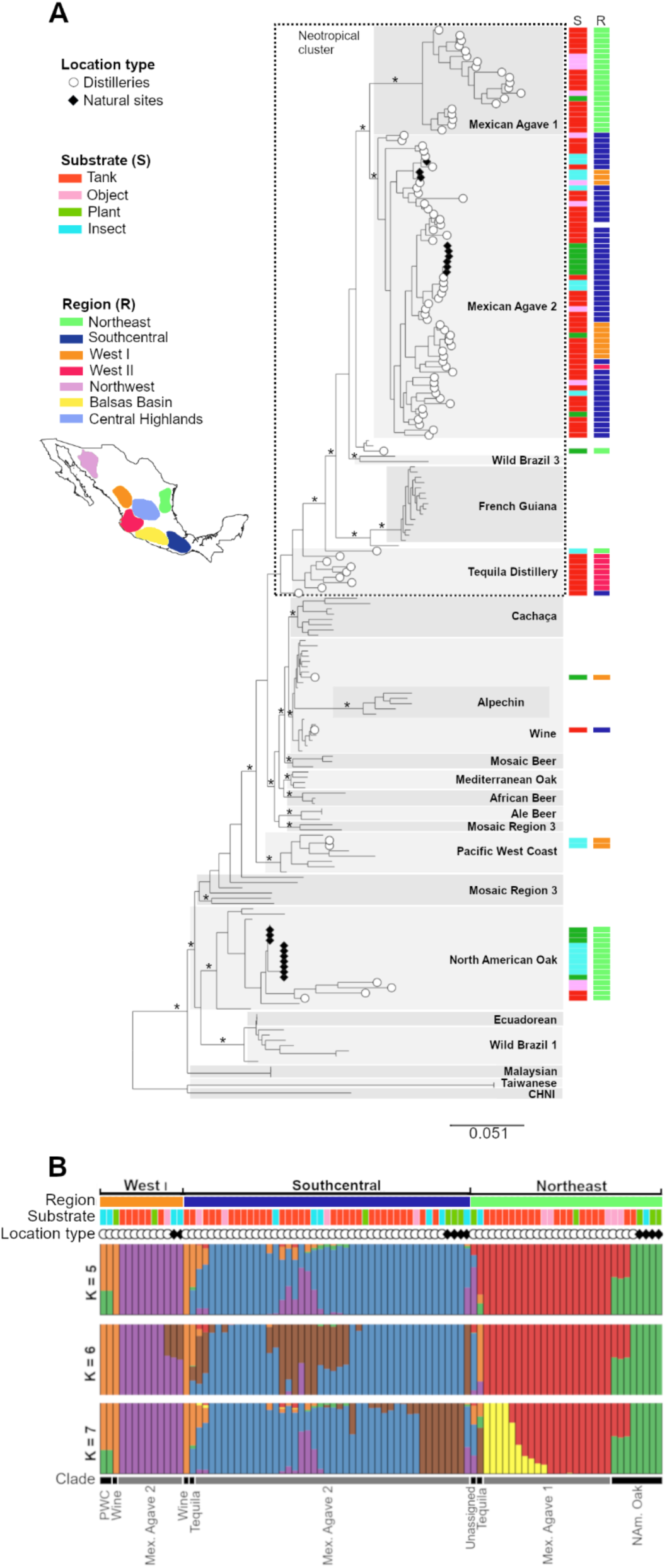
Relationships of *S. cerevisiae* strains from agave spirit distilleries and natural sites in Mexico. **(A)** Maximum likelihood tree using 870,190 biallelic SNPs of 203 strains. Strains from agave spirit distilleries include 42 isolates from this study, 56 from (Avelar-Rivas *et al*., 2024), and 8 from (Peter *et al*., 2018). All 19 *S. cerevisiae* isolates from natural sites in Mexico are from this study. The main clades of the world are shown with gray boxes following the nomenclature of (Peter *et al*., 2018) (with the exception of Wild Brazil 1 and 3 from (Barbosa *et al*., 2016) and Pacific West Coast from (Marr *et al*., 2023)); the Neotropical cluster was described by (Avelar-Rivas *et al*., 2024). Bootstrap support values ≥ 90 are shown with stars at the clade level and above (within-clade values are not shown for clarity purposes). Agave spirit producing regions include the three regions sampled in this study (Northeast, Southcentral and West I) and additional regions from which strains from Avelar-Rivas *et al*. (2024) were obtained from (Suppl. Table S1). **(B)** Population assignment analysis using ADMIXTURE. Region is shown at the top; location type is indicated with white circles (○, distilleries) and black diamonds (♦, natural sites). Substrate color code and clades correspond to those shown in (A). Assignation results for K = 2 to 15 are shown in Suppl. Mat. S3.

To assess the effect of geographical distribution on possible gene flow processes among sympatric populations, we ran ADMIXTURE on the genomes of the strains from Mexico (**Fig. 3B**). To avoid bias due to possible clonal replicates within the dataset (Alexander *et al*., 2009), clonal clusters were identified and a single strain per cluster was kept for the analysis. The analysis applied to a final set of 88 *S. cerevisiae* strains showed that in the West I and Southcentral regions the strains isolated from natural sites were not differentiated from those coming from distilleries. The additional genetic component at higher values of *K* that was associated to natural sites isolates (shown in brown) appears to reflect internal differentiation of the Mexican Agave 2 clade. This component was shared among strains from both the West I and Southcentral regions, and from both distilleries and natural sites. For example, at *K* = 5, the purple component was shared across regions and location types. At *K* = 6, the brown component was again shared between the two regions and across substrates. By *K* = 7, the brown component was restricted to the Southcentral region but still present in strains from both distilleries and natural sites.

The Northeast region also showed internal differentiation of distillery strains at *K* = 7, marked by a yellow component that includes six Mexican Agave strains from (Peter *et al*., 2018) and two strains from this study. Importantly, these strains also exhibited signs of admixture with the sympatric wild North American Oak clade (**Fig. 3B**). This evidence points to wild strains in the Northeast contributing to the genetic composition of strains found in the distilleries, but we did not find evidence of admixture processes in the opposite direction (from distilleries to wild strains). Additionally, we found no evidence of admixture among strains belonging to each of the two Mexican Agave groups. All the patterns previously described were stable at higher *K*, from *K* = 8 and up to *K* = 15 (**Suppl. Mat. S3**), with higher *K* values only revealing additional internal differentiation of Mexican Agave 1 and Mexican Agave 2 groups. Together, the phylogenetic and population structure analyses suggest that *S. cerevisiae* populations in natural sites and distilleries are largely the same, although no strains from the Mexican Agave 1 clade were found in natural environments.

### Three novel *S. paradoxus* SpB subgroups identified in Mexico

In contrast to *S. cerevisiae*, the population structure of *S. paradoxus* in Mexico and the broader Tropical Americas remains largely uncharacterized. To our knowledge, no genomic sequences of *S. paradoxus* strains from Mexico are currently available in public databases, and only two sequences exist from strains isolated in other Latin American countries, specifically in Brazil (Yue *et al*., 2017). As shown previously, MALDI-TOF identification indicated that *S. paradoxus* was more common in natural environments, with isolates of this species found in 13 of the 25 natural sites sampled. Notably, *S. paradoxus* coexisted with *S. cerevisiae* at 12 of these sites. In addition, *S. paradoxus* was isolated from six distilleries across all three sampled regions, as well as from agave fermentations in other producing areas. This was unexpected, given that *S. paradoxus* is typically considered rare in anthropogenic fermentative environments.

To understand the relationship of *S. paradoxus* populations from distilleries and natural environments we sequenced the genomes of 48 isolates from the three agave-spirit producing regions included in this study (11 from distilleries and 37 from natural sites), and 10 strains from agave fermentations from other distilleries in Mexico, collected by (Gallegos-Casillas *et al*., 2024) (**Table 1**). For the phylogenetic analyses we also included 52 strains from the recognized *S. paradoxus* clades of the world, for 110 strains in total (**Suppl. Table S2**) (Eberlein *et al*., 2019; Leducq *et al*., 2016; Xia *et al*., 2017; Yue *et al*., 2017). As shown in the phylogenetic tree (**Fig. 4A**), *S. paradoxus* isolates from Mexico clustered within the SpA and SpB clades. As expected, the majority (47 strains) belonged to SpB, which is considered native to North America. Of these, 20 strains were isolated from distilleries (13 from active fermentations and 7 from other substrates) and 27 from natural sites, including 3 from insects and 24 from plants.

**Fig. 4.**
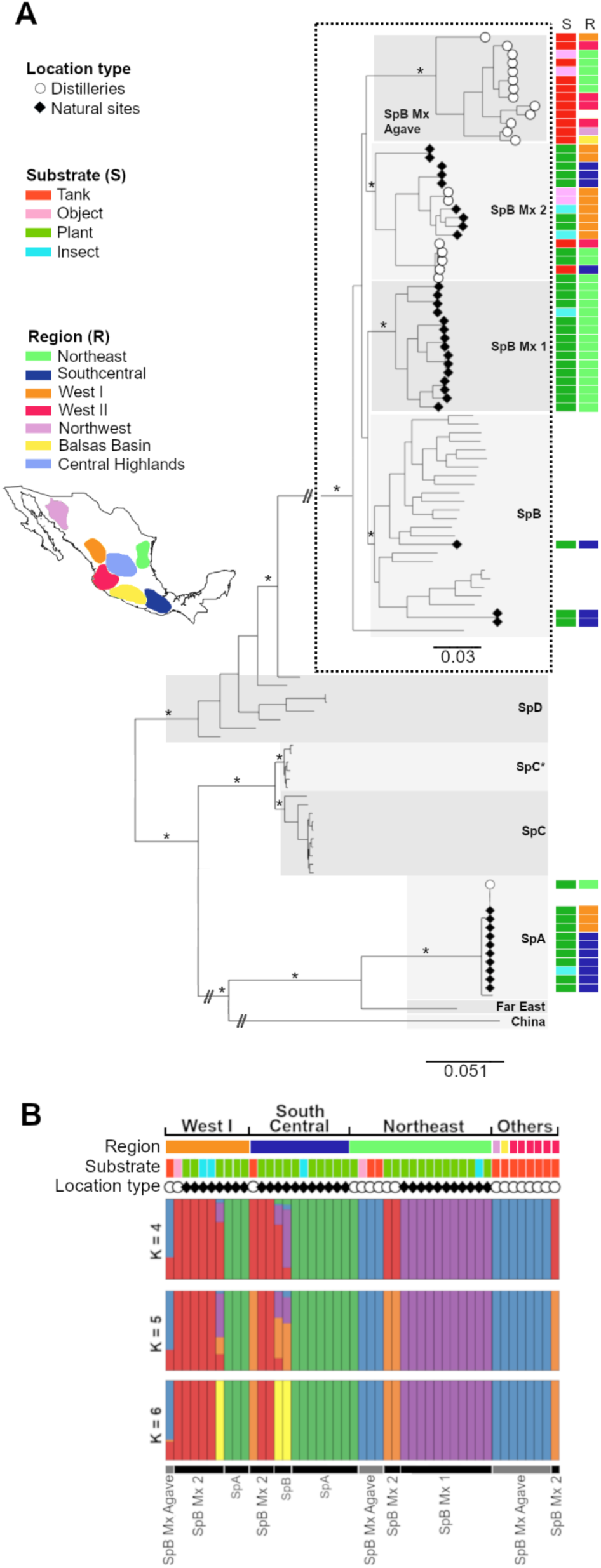
Relationships of *S. paradoxus* strains from agave spirit distilleries and natural sites in Mexico. **(A)** Maximum likelihood tree based on 876,358 SNPs from 110 strains. Strains from agave spirit distilleries (21 isolates) and natural sites (37 isolates) are from this study. The known clades of the species according to (Eberlein *et al*., 2019) and (He *et al*., 2022) are shown with gray boxes. Clade SpB has been slightly enlarged for clarity purposes (notice different scale of branch lengths). Bootstrap support values ≥ 90 are shown with stars at the clade level and above (within-clade values are not shown for clarity purposes). Agave spirit producing regions include the three regions sampled in this study (Northeast, Southcentral and West I) and additional regions from which strains from (Gallegos-Casillas *et al*., 2024) were obtained from (Suppl. Table S2). **(B)** Population assignment analysis using ADMIXTURE. Region is shown at the top, location type is signaled with white circles (○, distilleries) and black diamonds (♦, natural sites). Substrate color codes and clades correspond to those shown in A). Assignation results for *K* = 2 to 15 are shown in Suppl. Mat. S4.

The phylogeny revealed that the SpB isolates were actually structured into four subclades (**Fig. 4A**). One corresponds to the previously described SpB clade from the U.S.A. and Canada to which a few natural plant isolates from the Southcentral region were assigned. The remaining three subclades, all uncovered in this study, represent previously unrecognized populations that we herein name SpB_Mx. These include (i) SpB_Mx1, a large group of strains from the Northeast region, all isolated from natural sites; (ii) SpB_Mx2, which includes strains from both distilleries and natural sites; and (iii) SpB_MxAgave, composed exclusively of strains isolated from distilleries. We note that clade SpB_Mx1 is closely related to, but clearly distinct from, the North American SpB lineage and forms a sister group (**Fig. 4A**), also shown in purple in the ADMIXTURE plot (**Fig. 4B**). Subclades SpB_Mx2 and SpB_MxAgave form a monophyletic group that is more distantly related to the other SpB strains from Mexico, U.S.A., and Canada. In the ADMIXTURE results (**Fig. 4B**), these two groups are shown in red and blue, and both include strains from multiple agave-producing regions (**Fig. 4A**).

In addition, some *S. paradoxus* isolates were part of the SpA clade, with 11 strains from the three sampled geographical regions. The SpA lineage is thought to be native from Eurasia, from where it likely dispersed to North America “a sufficiently long time ago” to allow for divergence at microsatellite loci within the North American population (Kuehne *et al*., 2007). All SpA strains in this study came from natural areas in Mexico; in the West I region the strains were isolated in the core zone of a natural reserve that has minimal human presence. Given that the isolation sites are thousands of kilometers apart from each other, this pattern is consistent with the wide geographical distribution but low frequency of SpA in U.S.A. and Canada (Eberlein *et al*., 2019; Hyma & Fay, 2013; Sylvester *et al*., 2015). Our analysis also revealed that SpA and SpB strains coexist in the Southcentral and West I regions of Mexico. Notably, we isolated both lineages from a single oak bark sample collected at a natural site in the West I region (**Suppl. Table S2**). However, this coexistence was only observed in natural environments, as SpA strains were not associated with agave fermentations. Despite this coexistence, ADMIXTURE analyses suggested little to no genetic exchange between the SpA and SpB strains from Mexico (**Figure 4B**), as previously observed in North America (Kuehne *et al*., 2007). Indeed, SpA and SpB lineages remain differentiated from small to large *K* values, whereas at higher *K* values (e.g. *K* = 6) only internal differentiation of the SpB lineage was revealed (**Suppl. Mat. S4**). Reduced spore viability between SpA and SpB clades of North America suggests the existence of reproductive barriers that prevent genetic admixture between these sympatric populations (Charron *et al*., 2014b; Kuehne *et al*., 2007).

ADMIXTURE analyses also revealed that most coexisting *S. paradoxus* groups within each region appear to be genetically isolated from one another, showing no signs of admixture. The exceptions were two strains from the Southcentral region belonging to the SpB clade from USA and Canada and one strain from the West I region classified as SpB_Mx2, all of which showed evidence of admixture with other SpB groups (**Fig. 4B**; indicated by purple and orange at *K* = 5 and differentiated in yellow at *K* = 6). Altogether, our findings underscore that *S. paradoxus* populations in Mexico harbor substantial and previously unrecognized genetic diversity within the SpB lineage of the Americas, and that these lineages remain genetically distinct, even when occurring in sympatry.

### A previously unrecognized population of *S. paradoxus* persisting in agave fermentations

Our whole-genome analyses revealed distinct distributions of *Saccharomyces* spp. populations across natural and anthropogenic environments (**Fig. 5**). As previously described, *S. cerevisiae* was more frequent in distilleries, with Mexican Agave 1 and 2 clades dominating, although few strains from Mexican Agave 2 were also found in nearby natural sites. Wine and North American Oak clade strains occurred at lower frequencies in both environments. In contrast, *S. paradoxus* was more common in natural sites, where SpA, SpB, and SpB_Mx1 were virtually restricted to natural areas, while SpB_Mx2 occurred in both environment types.

**Fig. 5.**
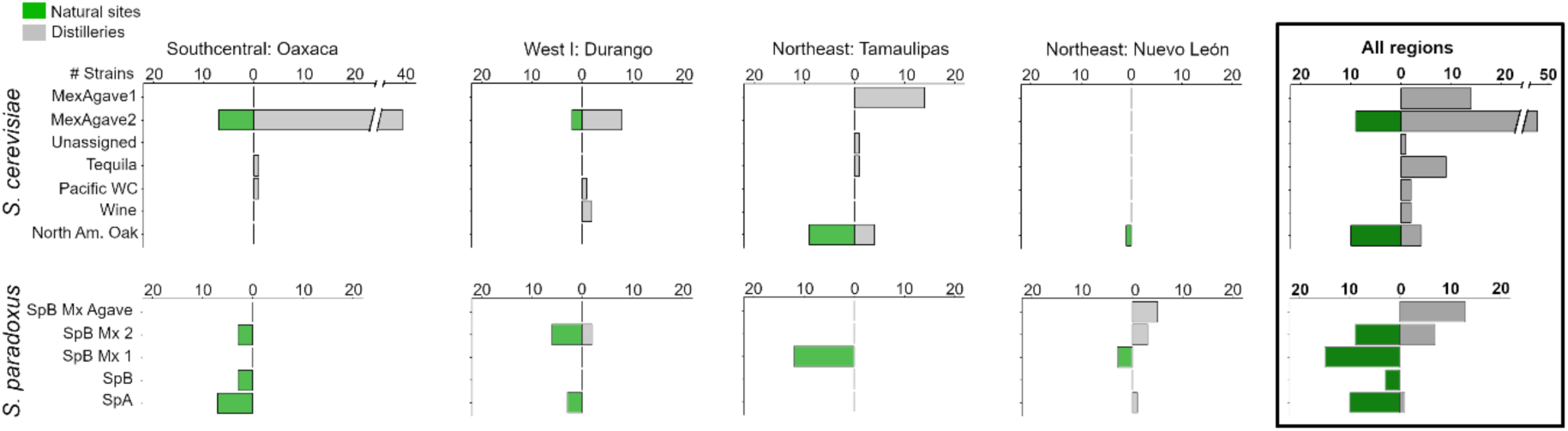
Clade distribution of *Saccharomyces* spp. among distilleries and natural sites in each of the three studied regions and across the country. The rightmost plots grouping all regions include strains isolated in this study and in (Avelar-Rivas *et al*., 2024; Gallegos-Casillas *et al*., 2024) (Suppl. Tables S1 and S2).

Unexpectedly, SpB_MxAgave was composed exclusively of strains isolated from distilleries (**Fig. 5**). Remarkably, this population, which is also the most genetically divergent from previously known SpB strains in North America, included 11 of the 13 *S. paradoxus* isolates obtained from active agave fermentation tanks. Strains belonging to this group were recovered from distilleries in five of the seven agave spirit-producing regions (Northeast, Northwest, West I, West II, and Balsas Basin), spanning up to 1,440 km. Notably, members of SpB_MxAgave were more genetically similar to each other than to other *S. paradoxus* strains isolated from different substrates within the same regions (**Fig. 4A**). However, no strains from this subclade were recovered from the state of Tamaulipas, the only producing region located east of the Sierra Madre Oriental mountain range, shown in **Figure 1C**.

The SpB_MxAgave was clearly differentiated in the ADMIXTURE analysis at different *K* values, as described above. Signatures of admixture with local wild strains were only found in a single strain from the West I region; the remaining SpB_MxAgave strains did not show any evidence of admixture with local wild SpB_Mx2 strains from the same region and even from the same exact location (**Fig. 4B**). In contrast, strains belonging to SpB_Mx2 that come from plants in distilleries grouped with other strains from the same geographical region and have the corresponding genetic components in the ADMIXTURE analysis (**Fig. 4B**). These results suggest that traditional agave fermentations represent a distinct ecological habitat for a specific *S. paradoxus* population, potentially driving differentiation within the species. The SpB_MxAgave population may therefore be undergoing adaptation to an anthropogenic environment, which, to our knowledge, has not previously been reported for the yeast *S. paradoxus*. Taken together, our findings uncover an unrecognized reservoir of genetic and ecological diversity in *S. paradoxus* and highlight agave fermentations as a unique system for studying evolution of the species at the wild-anthropogenic interface.

## Discussion

*Saccharomyces* yeasts are emerging as powerful models for understanding the ecological factors that shape microbial eukaryotic populations and the evolutionary processes underlying adaptive divergence during domestication (De Chiara *et al*., 2022; Replansky *et al*., 2008; Tengölics *et al*., 2024; Yue *et al*., 2017). However, most ecological genomics studies of *S. cerevisiae* and *S. paradoxus* have been conducted in temperate regions (Eberlein *et al*., 2019; Peris *et al*., 2023), with notable exceptions from Brazil (Barbosa *et al*., 2016; Barbosa *et al*., 2018), where the two species have been shown to coexist and interbreed. Here, through extensive field sampling and genome-wide analyses, we provide new insights into the ecology, distribution, and population structure of *S. cerevisiae* and *S. paradoxus* in traditional agave fermentations in Mexico. Our findings advance the understanding of yeast biogeography in tropical regions and highlight traditional agave fermentations as dynamic systems for investigating microbial adaptation at the interface between wild and human-influenced environments.

### *S. cerevisiae* and *S. paradoxus* coexist in Mexico but differ in habitat association

We observed clear differences in habitat preference between the two yeast sister species. While *S. cerevisiae* was more abundant in distilleries, *S. paradoxus* was predominantly isolated from natural environments. These results align with previous ecological characterizations of these species (Liti *et al*., 2009; Sniegowski *et al*., 2002). Nonetheless, we also found *S. cerevisiae* in natural sites and, more unexpectedly, *S. paradoxus* within distilleries, including strains isolated directly from fermentation tanks. This finding challenges the strict interpretation of *S. paradoxus* as a purely wild species and suggests that, under certain conditions, it may persist in anthropogenic fermentative environments. It is worth noting that the general absence of *S. paradoxus* from most human-made fermentations remains puzzling, especially given its ability to perform relatively well in wine and beer fermentations (Dashko *et al*., 2016; Nikulin *et al*., 2020).

Moreover, our data did not reveal a clear relationship between the differential distributions of the species and climatic conditions such as mean annual temperature (**Suppl. Mat. S5**). This contrasts with temperate-region studies showing clear temperature-based distributions (Gonçalves *et al*., 2011; Charron *et al*., 2014a; Leducq *et al*., 2014; Robinson *et al*., 2016), highlighting that temperature alone does not fully explain species distribution patterns observed in tropical regions. Future studies focusing on growth temperature optima and fitness of *Saccharomyces* from the tropics under diverse conditions will further contribute to clarifying these ecological relationships.

### *Saccharomyces* yeasts in tropical Mexico are associated with diverse plant hosts beyond oaks

In temperate regions, oak trees (*Quercus*) have been described as primary natural habitats for *Saccharomyces* species (Hyma & Fay, 2013; Naumov *et al*., 1998; Peris *et al*., 2023; Sampaio & Gonçalves, 2008; Sniegowski *et al*., 2002; Wang *et al*., 2012). However, despite Mexico being a center of oak diversity (Kremer & Hipp, 2020), our results showed that *Saccharomyces* spp. associated similarly with multiple plant families, including Anacardiaceae and Fabaceae. Specifically, the isolation rates from oaks (12%) were comparable to those from other plant families (8–14%), suggesting that plant hosts in tropical regions may be more diverse than previously acknowledged. This broader host range in tropical ecosystems points to possible overlooked ecological interactions. For instance, Anacardiaceae and Fabaceae plants are both known for their extrafloral nectaries, which secrete nutrient-rich exudates potentially utilized by yeasts (Weber & Keeler, 2013). Given the high plant species turnover in tropical areas (McFadden *et al*., 2019), systematically investigating yeast associations with diverse plant families could reveal additional hidden diversity in *Saccharomyces*.

### Insects may mediate yeast exchange between natural and anthropogenic habitats in Mexico

In our study, *Saccharomyces* yeasts were isolated from insects, suggesting they play a role in facilitating the dispersal of both *S. cerevisiae* and *S. paradoxus*. Isolation rates from Drosophilidae were highest compared to other insect families, consistent with their known association with yeasts (Meriggi *et al*., 2020). Interestingly, insect-associated strains within distilleries grouped closely with strains from fermentation tanks and objects, further supporting their possible role as vectors within anthropogenic habitats, as previously suggested (Lachance, 1995). In contrast, isolation rates from insects were considerably lower in natural areas, where only Drosophilidae yielded isolates from both yeast species. This difference suggests that insect-mediated dispersal is complex and influenced by habitat-specific ecological factors and insect physiology, which requires further research to clarify the underlying mechanisms.

### A distillery-associated *S. paradoxus* population suggests adaptation to anthropogenic habitats

Unexpectedly, among the previously unrecognized populations within the SpB clade, we identified SpB_MxAgave, a *S. paradoxus* population exclusively associated with distillery fermentation tanks. While isolated cases of *S. paradoxus* have previously been reported in fermentation environments (Dashko *et al*., 2016; Gallegos-Casillas *et al*., 2024), they have been rare and inconsistent, despite the species’ fermentative capacity (Nikulin *et al*., 2020). To our knowledge, SpB_MxAgave represents the first clear example of a genetically distinct *S. paradoxus* lineage consistently associated with an anthropogenic habitat. This population was geographically widespread across multiple agave-producing regions, spanning distances of up to 1,440 km, yet remained genetically distinct without evidence of admixture with local wild populations. Moreover, the coexistence of distillery-exclusive populations in both *S. cerevisiae* (Mexican Agave 1) and *S. paradoxus* (SpB_MxAgave) suggests convergent adaptation to agave fermentations, which constitute a unique fermentative environment (Colón-González *et al*., 2025). These discoveries highlight traditional agave fermentations as unique systems for studying microbial adaptation processes in human-associated habitats.

### Genetic admixture occurs among sympatric yeast populations

Our genome-wide analyses revealed admixture events between wild and domesticated lineages, such as gene flow from introduced strains from the Wine clade into local Mexican Agave 2 populations of *S. cerevisiae*. Similarly, North American Oak clade strains isolated from distilleries exhibited evidence of admixture with local domesticated populations, suggesting active interpopulation dynamics rather than spontaneous colonization into fermentative environments. In contrast, despite geographical overlap, we observed limited admixture between major *S. paradoxus* lineages (SpA and SpB), consistent with previously documented reproductive isolation (Charron *et al*., 2014b). These findings highlight that divergent lineages can coexist at small spatial scales (Hyma & Fay, 2013; Koufopanou *et al*., 2006; Sylvester *et al*., 2015; Xia *et al*., 2017), yet their genetic interactions remain limited and are shaped by broader ecological contexts.

### Concluding remarks

Overall, our study expands current knowledge of yeast ecology and biogeography, particularly in the understudied tropical regions of the Americas. Beyond their ecological significance, traditional agave fermentations emerge as a powerful natural laboratory for exploring the genetics and evolution of *Saccharomyces* species. The coexistence, diversification, and potential adaptation of yeast populations in this system offer unique opportunities to investigate microbial ecology at the interface of wild and anthropogenic environments. Moreover, the discovery of several previously undescribed populations within the SpB clade of *S. paradoxus* highlights an important genetic reservoir for this species. These insights are not only essential for understanding the evolutionary dynamics shaping yeast communities but also have important implications for the conservation of traditional agave fermentations. These practices are deeply rooted in cultural heritage and rely on microbial populations sustained by local biodiversity and traditional production methods. Ensuring the long-term sustainability of agave spirit production will require protecting surrounding natural habitats and preserving artisanal fermentation practices in Mexico and globally.

## Materials and Methods

### Sampling and locations

Sampling was performed during the year 2021, in three regions: Southcentral (Oaxaca state), West I (Durango state) and Northeast (Nuevo León and Tamaulipas states). Sampling areas are representative of different biogeographic regions: Oaxaca state is found in the *Sierra Madre del Sur* province, Durango is part of the *Chihuahuan Desert* province, Nuevo León is in the *Sierra Madre Oriental*, and Tamaulipas in the *Tamaulipas* province (Morrone *et al*., 2017). 15 distilleries and 26 natural sites were visited. Most distilleries had been previously sampled by the Yeast Genomes MX consortium (Gallegos-Casillas *et al*., 2024).

Natural sites were defined as locations situated at least ∼3 km and up to 42 km from sampled distilleries, where human presence (e.g., constructions, agricultural fields, roads) was minimal and where oak trees (genus *Quercus*, Fagaceae) were present; yet, other trees in the transect belonging to different plant families were also sampled. The average distance of the natural site from the closest distillery was 9.2 km; only one natural sites in the Northeast region was located 1 km away, but this was in a rural area with very little human presence.

#### Fermentation tanks

To collect samples from fermentation tanks in distilleries, an aliquot was taken with a sterile serological pipette at arm’s reach and four ml were stored in a sterile cryovial with glycerol to a final concentration of 25%, as previously described (Gallegos-Casillas, 2024). Two other cryovials were also filled but without glycerol. All cryovials were stored in liquid nitrogen until arrival at the laboratory, where they were stored at -70°C until further processing.

#### Objects

In active distilleries, recently cooked agave stems, waste of agave grinding, solid remains of previous fermentations in the tanks, firewood, pounders, and other wood and metal tools were collected. When distilleries were closed or without production at that moment, sampling from tools, agave residues, wood, and other objects was performed. Depending on the object sampled, different tools were used for collection; all tools were previously sterilized with 10% chlorine first and 70% ethanol afterwards.

#### Insects

Insects were actively trapped with an entomological net previously sterilized in the laboratory with UV radiation for two hours in a laminar flow hood. The net was swiped 1 m above the ground until 10-20 insects were trapped, then put in a conical sterile 50 mL tube with an autoclaved custom-made aspirator. Within a single location, after each use, nets were sterilized by soaking them in 70% ethanol. When moving to a new location, a sterile net previously UV sterilized was used to avoid horizontal contamination. Insects were identified at the level of Order in the field and in some cases at the level of Family. Afterwards, the 50 mL tubes were stored in a cooler with ice until further processing in the laboratory.

#### Plants

In distilleries, bark from trees growing within or at the border of the facilities with a minimum trunk diameter at breast height (DAB) of 10 cm was collected. In natural sites, 50 x 10 m transects were defined, where bark of trees with DAB > 10 cm was sampled, including oaks and other tree species present in the transect. Eight to 20 g of bark was collected in 50 mL sterile conical tubes with a cork borer or knife and a tweezer. The borer, knife, and tweezers were autoclaved before sampling and serially cleaned with 10% chlorine followed by 70% ethanol solutions between each sample to avoid horizontal contamination. The solutions of ethanol and chlorine were stored in 50 mL aliquots in conical tubes where the tool was submerged for 1 min before use, and new aliquots were used at each location. Samples collected in the conical tubes were stored in a cooler with ice for 3-5 days until further processing in the laboratory. Photos of the leaves, bark, flowers, and fruits, if present, were taken. For each morphospecies in a single location, a branch with leaves (and, if present, flowers or fruits) was collected and preserved in a botanical press for taxonomic identification.

#### Data collection of field samples

Sample metadata (date, location type, name of location, geographical coordinates, type of sample, photograph, name of collector) were recorded on the KoboToolbox server using the KoboCollect application. Templates were created in the Kobo server, which were filled for each sample in the field using the corresponding mobile app. Data cleaning was performed in *python* v3.8.8 and *OpenRefine* v3.4.1.

### Taxonomic identification of isolates

Yeast enrichment in the laboratory was performed with the protocol of (Liti *et al*., 2017) with modifications. The 50 mL conical tube containing the sample was directly used for enrichment. 20 mL of Saccharomyces Sensu Stricto Enrichment Medium were added and incubated for up to three weeks at 25°C regularly inspecting them for signs of fermentation, like sedimentation, turbidity, and CO_2_ production. The actively fermenting cultures were diluted 10^-3^ or 10^-4^ depending on the level of saturation and plated using glass beads in petri dishes with 20 mL of WL Nutrient Agar (SIGMA and Difco). This medium allows morphological differentiation of yeast colonies due to their pH indicators. Petri dishes were incubated at 25°C for 2-5 days until colonies were well defined, and the agar medium changed from blue to green and then refrigerated at 4°C until strains selection. For strain selection, 12 colonies with *Saccharomyces*-like morphology and 12 with varied morphologies were selected and placed in 150 uL of liquid YPD medium in a 96 well plate. Plates were incubated at 30°C for two days, 100 uL of 50% glycerol was then added and strains were stored at -70°C.

For taxonomic identification of isolates, the protocol of (Avelar-Rivas *et al*., 2024) and (Gallegos-Casillas *et al*., 2024) was used using matrix-assisted laser desorption ionization–time of flight mass spectrometry (MALDI-TOF) with Biotyper® that determines the unique proteomic fingerprint of an organism and matches its characteristic mass spectrum pattern with an extensive reference library to determine the organism identity at the species level. The output of the instrument is a score that ranges from 0 to 3 and that reflects whether the tested isolate belongs to a given species in the reference database. Isolates with a score below 1.5 cannot be identified reliably. Direct bulk biomass was assessed from colonies growth in solid YPD medium. In some cases, protein extraction was carried out to improve the quality of the mass spectrum.

Overall, 861 samples were collected in the field that translated into 4,006 yeast isolates. From all isolates, 1,094 isolates (from 139 samples) were identified to be *Saccharomyces* with MALDI-TOF. Strains were selected for genome sequencing based on the list of ten ranked probable identities according to their score value from the MALDI-TOF, geographical origin and substrate from which they were isolated.

### Microbial community statistics

Taxonomic identities of isolates with MALDI-TOF score >1.5 were used to estimate species diversity with Shannon’s diversity index (*H’*) and Simpson’s heterogeneity index (*D*) per location and substrate type (library *tidyverse* in *R* v4.4.2; (Wickham *et al*., 2019)). Strains isolated at each location type were grouped and Fisher’s exact tests were performed to test if the number of strains of *S. cerevisiae* and *S. paradoxus* was higher than expected by random in distilleries and natural sites (in *R* v4.2.2).

### DNA extraction and genome sequencing

DNA extraction was performed using a protocol modified from the MasterPure^TM^ Yeast DNA Purification Kit. In brief, a lysis buffer for enzymatic lysis with zymolyase, lyticase and RNAse A was used, and precipitated with isopropanol cooled at -20°C. DNA yield was quantified with a Qubit^TM^ fluorescence spectrophotometer, and the 260/280 and 260/230 ratios were determined with a ND-1000 NanoDrop^®^. Short-read DNA sequencing library preparation and sequencing was performed at the BGI using the DNBSeq platform with 150 bp paired-end reads for each strain.

### Quality control and genome mapping

BGI reported a quality assessment of the raw reads with *SOAPnuke*, which indicated high overall sequencing quality. Adapters and low-quality reads were further removed with *fastp* v0.20.0 (Chen *et al*., 2018). Afterwards, raw reads were mapped to a concatenated genome of the eight species of the *Saccharomyces* genus, plus the reference genomes of *Kluyveromyces marxianus* and *Pichia kudriavzevii*, with *bwa* v0.7.4 (Li & Durbin, 2009). The references genomes were the following: *S. cerevisiae*, we used the version R64-2-1_20150113 downloaded from the SGD, accessed Jan 16 2020; *S. paradoxus* YPS138 was downloaded from the Yeast Population Reference Panel (YPRP) https://yjx1217.github.io/Yeast_PacBio_2016/data/, accessed Feb 19 2020; *S. jurei* GCA_900290405.1, *S. kudriavzevii* GCA_900682695.1; *S. arboricola* GCF_000292725.1; *S. eubayanus* GCA_001298625.1; *S. mikatae* IFO 1815^T^ and *S. uvarum* CBS 7001 were downloaded from https://sss.genetics.wisc.edu (Scannell *et al*., 2011), accessed Feb 2022; *K. marxianus* GCF_001417885.1; *P. kudriavzevii* GCF_003054445.1.

Plots with the coverage depth to the concatenated reference genomes were made using *samtools* v1.9 and *R* v3.6.1. A custom python script was used to assign species identity if > 90% of the reads were mapped to a given species.

### Variant calling and genotyping

Variant calling was performed with *GATK* v4.1.1.0 (DePristo *et al*., 2011) from the read alignments to the eight reference genome sequences following the best practice workflow manual for quality control of SNP calling, and 10% of missing SNPs per sample was allowed. Strains that were identified by genome coverage as *S. cerevisiae* were extracted and mapped to the S288C reference strain. During variant calling we also included 40 strains from agave fermentation tanks isolated from other Mexican regions (Avelar-Rivas *et al*., 2024), 5 strains isolated by Lachance (Lachance, 1995) sequenced by (Avelar-Rivas *et al*., 2024) and 105 strains representative of other world clades (Peter *et al*., 2018) (**Suppl. Table S1**). The final matrix contained 203 *S. cerevisiae* genomes and 870,190 biallelic SNPs.

A similar matrix of SNPs was generated with 107 *S. paradoxus* strains mapped to reference strain YPS138. Strains were distributed as follows: 48 from the three sampled regions (West I, Northeast, Southcentral), 10 from previous collection efforts in these and other producing regions of Mexico (previously not published and made available in this study; (Gallegos-Casillas *et al*., 2024), and 50 representatives of the recognized lineages of the species SpA, SpB, SpC, SpC*, and SpD (Eberlein *et al*., 2019; He *et al*., 2022; Xia *et al*., 2017; Yue *et al*., 2017) (**Suppl. Table S2**). The final matrix included 110 strains and 876,358 biallelic SNPs.

### Phylogenetic reconstruction of the sequenced strains

The matrixes with the genotyped SNPs were aligned and converted from the default *GATK* vcf format to phy format with a python script (Ortiz, 2019). Afterwards, the phylogenetic trees for each species, *S. cerevisiae* and *S. paradoxus*, were built using the maximum likelihood algorithm of *RAxML* v8.2.12 and a GTR substitution model (Stamatakis, 2014). We used 100 bootstraps to assess branch support.

### Population admixture analysis

Population analysis was performed using *ADMIXTURE* v1.3.0 (Alexander *et al*., 2009). To avoid any bias in the analysis related to possible clonal replicates, strains that were isolated from the same sample and that displayed a phylogenetic distance of < 0.005 in the ML phylogenetic tree were removed from the analysis. In those cases, one strain from the group of isolates suspected to be clones was chosen randomly. The final ADMIXTURE analysis of *S. cerevisiae* was run on 88 strains (36 from this study, 46 from (Avelar-Rivas *et al*., 2024), and 6 from the Mexican Agave clade of (Peter *et al*., 2018)). The same SNPs matrix used for the phylogeny was filtered to keep these samples and pruned for linkage disequilibrium (LD) with *plink* v1.9 (`--indep-pairwise 50 5 0.5`), keeping a final number of 108,636 SNPs. We performed the ADMIXTURE analysis of *S. paradoxus* in the same way, keeping a final set of 47 strains (all from this study) and 86,448 SNPs. ADMIXTURE was run on the output bed matrix for K1-K15 and 10 seeds. The best K values were determined according to the lowest cross-validation error values (**Suppl. Mat. S3 and S4**).

## Data availability

The raw sequence reads of 100 *Saccharomyces* strains sequenced here are under SRA project PRJNA1232640.

## Acknowledgements

We are grateful to Oscar Estrada, Gerardo Hernández, and Gilberto Roldán (Durango), Cuauhtémoc Jacques (Tamaulipas), and Karina Abad, Graciela Ángeles and Edgar Ángeles (Oaxaca) for their invaluable assistance during the field work described in this study. We thank Luis Aguilar, Vanessa Arellano, Antonio Basurto, Marco Chavez, Mauricio Campa, Alejandra Castillo, Maritrini Colón, Porfirio Gallegos, Jair García, Luis F. García, Antonio Hernández, Martín Moreno, Susana Ruiz, Carina Uribe, and Karen Vaca for technical assistance, and Anne Gschaedler and Marc Andre Lachance for sharing their strains.

## SUPPLEMENTARY MATERIALS

Ecological and genomic divergence of *Saccharomyces* species across wild and fermentative environments in tropical Mexico

López, Aguirre-Dugua, et al. 2025

**Supplementary Material S1 –** Climatic conditions and photographs of sampled locations

**Supplementary Material S2 –** Diversity of microbial species isolated with the enrichment medium per location type and substrate.

**Supplementary Material S3 –** ADMIXTURE analyses of *Saccharomyces cerevisiae* strains from agave-spirits distilleries and natural sites of Mexico

**Supplementary Material S4 –** ADMIXTURE analyses of *Saccharomyces paradoxus* strains from agave-spirits distilleries and natural sites of Mexico

**Supplementary Material S5 –** Presence of *Saccharomyces* species in distilleries and natural sites according to climatic conditions of sampled localities

## SUPPLEMENTARY MATERIAL S1

**Climatic conditions and photographs of sampled locations**

**Suppl. Mat. S1.** Climatic characteristics of sampled localities (based on WorldClim data; (Fick & Hijmans, 2017).

| Location ID | Region | State | Location Type | Mean Annual Temp (°C) | Maximum Temperature Warmest Month (°C) | Mean Annual Precip (mm) | Elevation (masl) |
| --- | --- | --- | --- | --- | --- | --- | --- |
| 1 | Southcentral | Oaxaca | Distillery | 20.86 | 29.10 | 672 | 1667 |
| 7 | Southcentral | Oaxaca | Distillery | 19.83 | 27.39 | 647 | 1768 |
| 13 | Southcentral | Oaxaca | Distillery | 21.36 | 29.60 | 682 | 1600 |
| 26 | Northeast | Tamaulipas | Distillery | 21.95 | 34.40 | 707 | 586 |
| 44 | West I | Durango | Distillery | 18.26 | 32.09 | 452 | 1757 |
| 45 | West I | Durango | Distillery | 18.21 | 32.09 | 452 | 1740 |
| 48 | West I | Durango | Distillery | 18.11 | 31.89 | 462 | 1783 |
| 85 | Southcentral | Oaxaca | NaturalEnv | 20.87 | 29.10 | 683 | 1653 |
| 86 | Southcentral | Oaxaca | NaturalEnv | 20.87 | 29.10 | 683 | 1653 |
| 87 | Southcentral | Oaxaca | NaturalEnv | 19.82 | 27.39 | 641 | 1751 |
| 88 | Southcentral | Oaxaca | NaturalEnv | 19.09 | 26.29 | 632 | 1871 |
| 89 | Southcentral | Oaxaca | NaturalEnv | 18.4 | 25.39 | 669 | 1865 |
| 90 | Southcentral | Oaxaca | NaturalEnv | 19.82 | 27.39 | 641 | 1751 |
| 91 | Southcentral | Oaxaca | NaturalEnv | 19.26 | 26.5 | 627 | 1890 |
| 92 | Southcentral | Oaxaca | NaturalEnv | 19.09 | 26.29 | 632 | 1871 |
| 93 | Southcentral | Oaxaca | NaturalEnv | 18.88 | 26.10 | 656 | 1836 |
| 94 | Southcentral | Oaxaca | NaturalEnv | 19.26 | 26.5 | 627 | 1890 |
| 95 | Southcentral | Oaxaca | NaturalEnv | 19.26 | 26.5 | 627 | 1890 |
| 96 | West I | Durango | NaturalEnv | 11.91 | 22.89 | 759 | 2799 |
| 97 | West I | Durango | NaturalEnv | 11.91 | 22.89 | 759 | 2799 |
| 98 | West I | Durango | NaturalEnv | 12.01 | 23 | 753 | 2809 |
| 100 | West I | Durango | Distillery | 18.21 | 32.09 | 452 | 1740 |
| 101 | West I | Durango | NaturalEnv | 17.30 | 30.39 | 461 | 1926 |
| 102 | West I | Durango | NaturalEnv | 16.79 | 29.29 | 484 | 2014 |
| 103 | Northeast | Tamaulipas | NaturalEnv | 22.06 | 34.40 | 732 | 605 |
| 104 | Northeast | Tamaulipas | NaturalEnv | 21.43 | 32.90 | 756 | 1082 |
| 105 | Northeast | Tamaulipas | NaturalEnv | 21.43 | 32.90 | 756 | 1082 |
| 107 | Northeast | Tamaulipas | NaturalEnv | 21.43 | 32.90 | 756 | 1082 |
| 108 | Northeast | Tamaulipas | NaturalEnv | 21.43 | 32.90 | 756 | 1082 |
| 109 | Northeast | Tamaulipas | Distillery | 22.04 | 34.09 | 704 | 591 |
| 110 | Northeast | Tamaulipas | Distillery | 22.38 | 35 | 739 | 468 |
| 111 | Northeast | Tamaulipas | Distillery | 21.88 | 33.90 | 705 | 628 |
| 112 | Northeast | Tamaulipas | Distillery | 22.31 | 34.90 | 735 | 500 |
| 113 | Northeast | Tamaulipas | NaturalEnv | 21.68 | 33.5 | 703 | 792 |
| 114 | Northeast | Tamaulipas | Distillery | 23.05 | 35.70 | 714 | 358 |
| 115 | Northeast | Tamaulipas | NaturalEnv | 16.31 | 28.29 | 412 | 2092 |
| 116 | Northeast | Nuevo Leon | Distillery | 13.95 | 23.89 | 595 | 2661 |
| 117 | Northeast | Tamaulipas | NaturalEnv | 16.47 | 28.5 | 410 | 2070 |
| 118 | Northeast | Nuevo Leon | NaturalEnv | 14.16 | 24.29 | 570 | 2506 |
| 119 | Northeast | Nuevo Leon | Distillery | 13.95 | 23.89 | 595 | 2661 |

**Suppl. Mat. S1.B.**
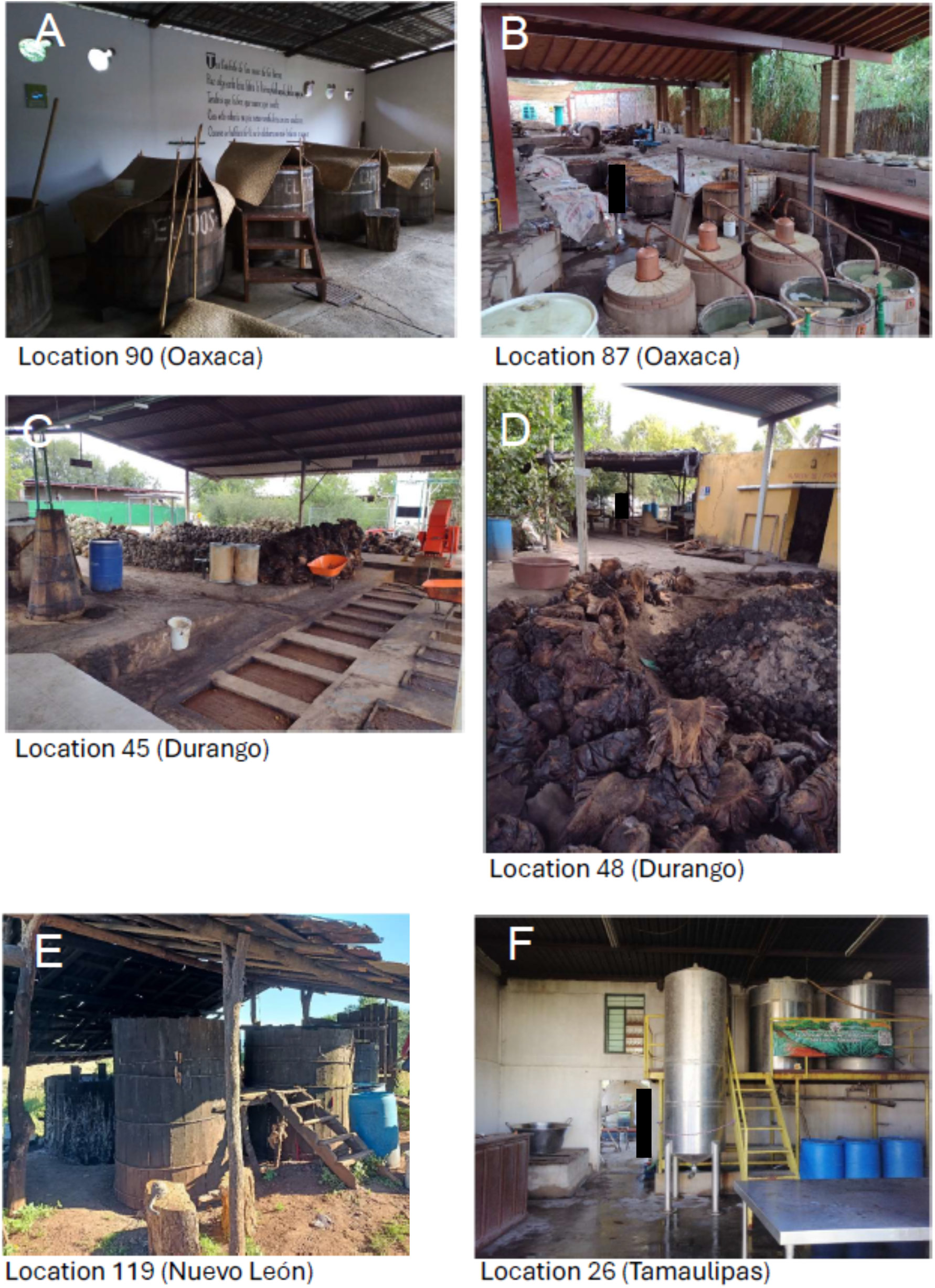
Photographs of representative traditional distilleries from the three regions (A-B: Southcentral; C-D: West I; E-F: Northeast) (black rectangles cover individuals that are not authors).

**Suppl. Mat. S1.C.**
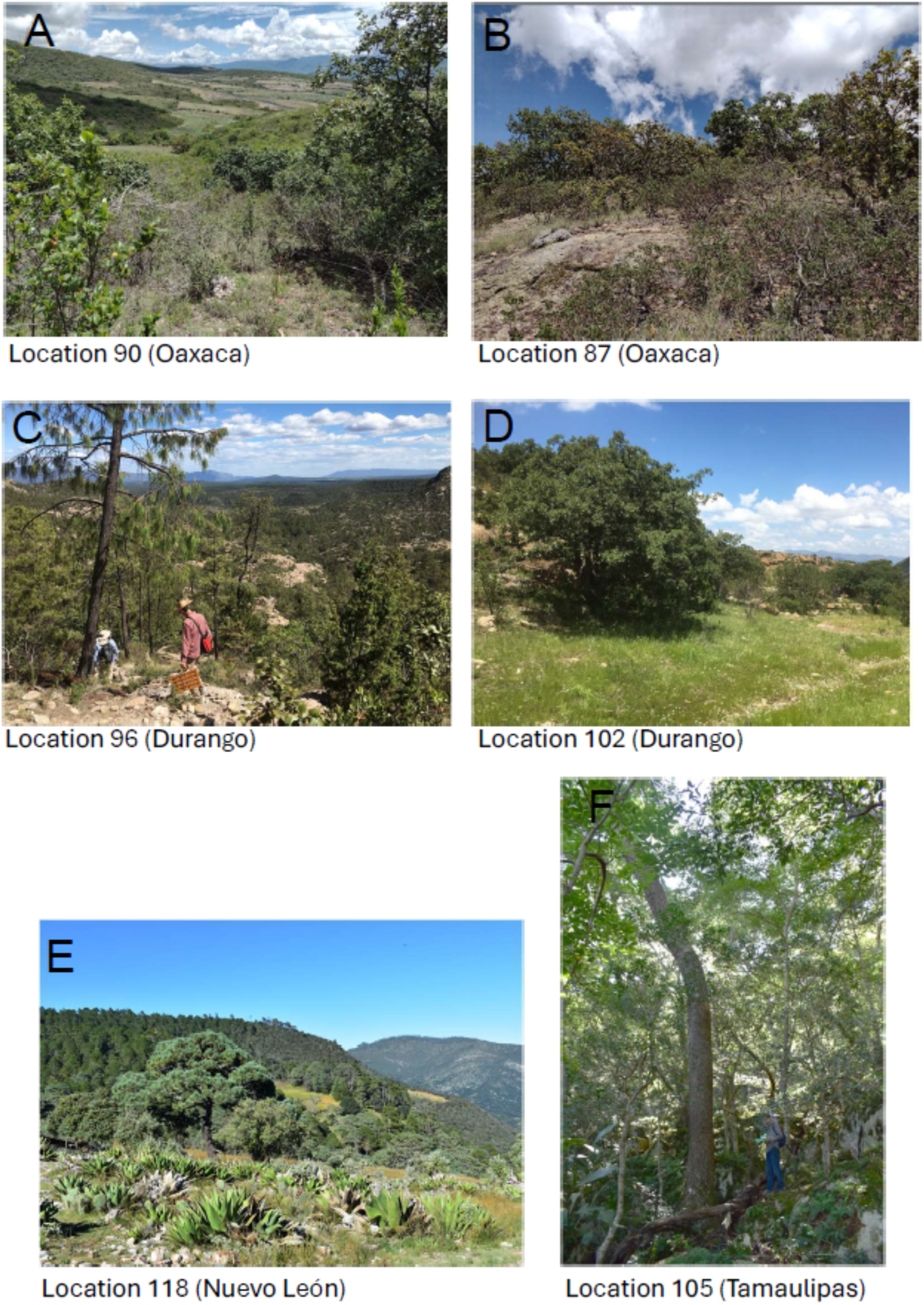
Photographs of representative natural sites sampled in this study (A-B: Southcentral; C-D: West I; E-F: Northeast) (shown individuals are the authors of the work).

## SUPPLEMENTARY MATERIAL S2

**Diversity of microbial species isolated with the enrichment medium per location type and substrate**

**Suppl. Mat. S2.A.**
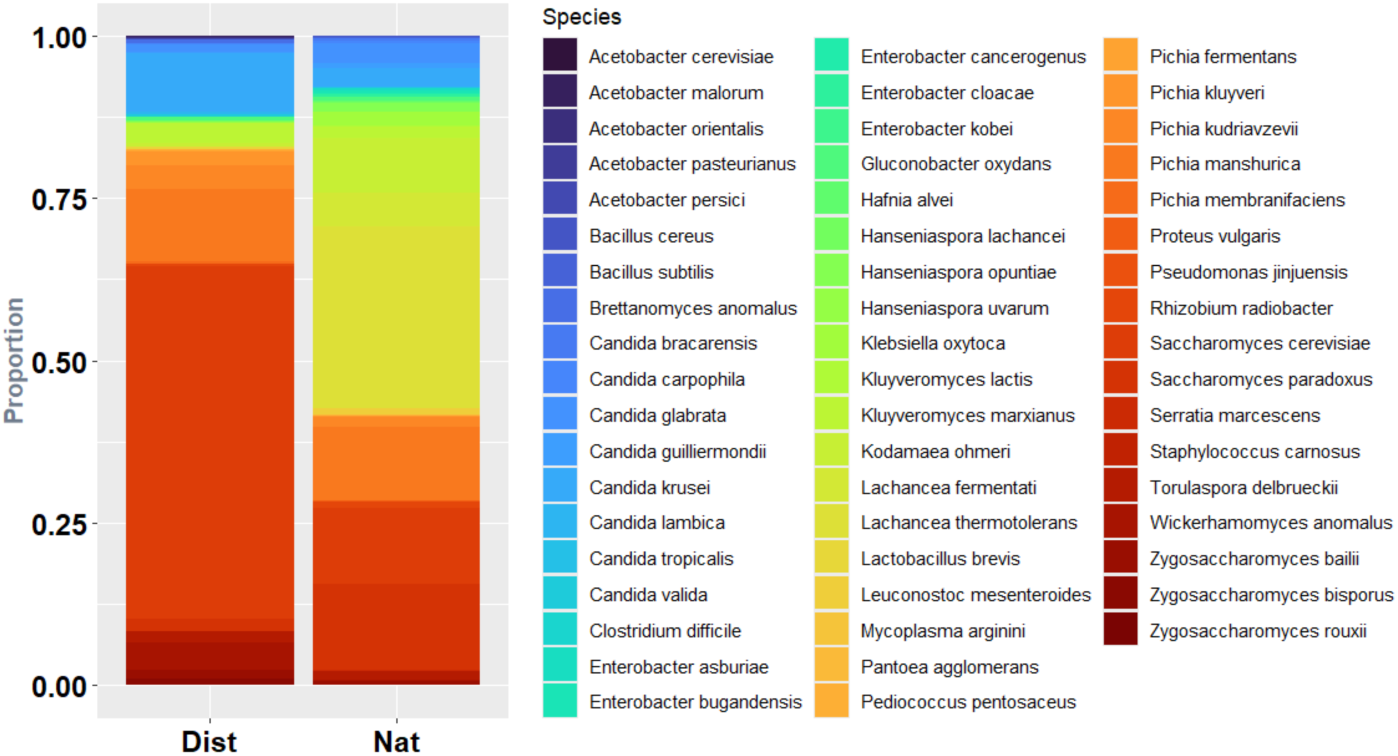
Diversity of species isolated in both location types and confidently identified with MALDI-TOF (score > 1.5). *Dist*: distilleries (*n* = 1,375 isolates). *Nat*: Natural sites (*n* = 834 isolates).

**Suppl. Mat. S2.B.**
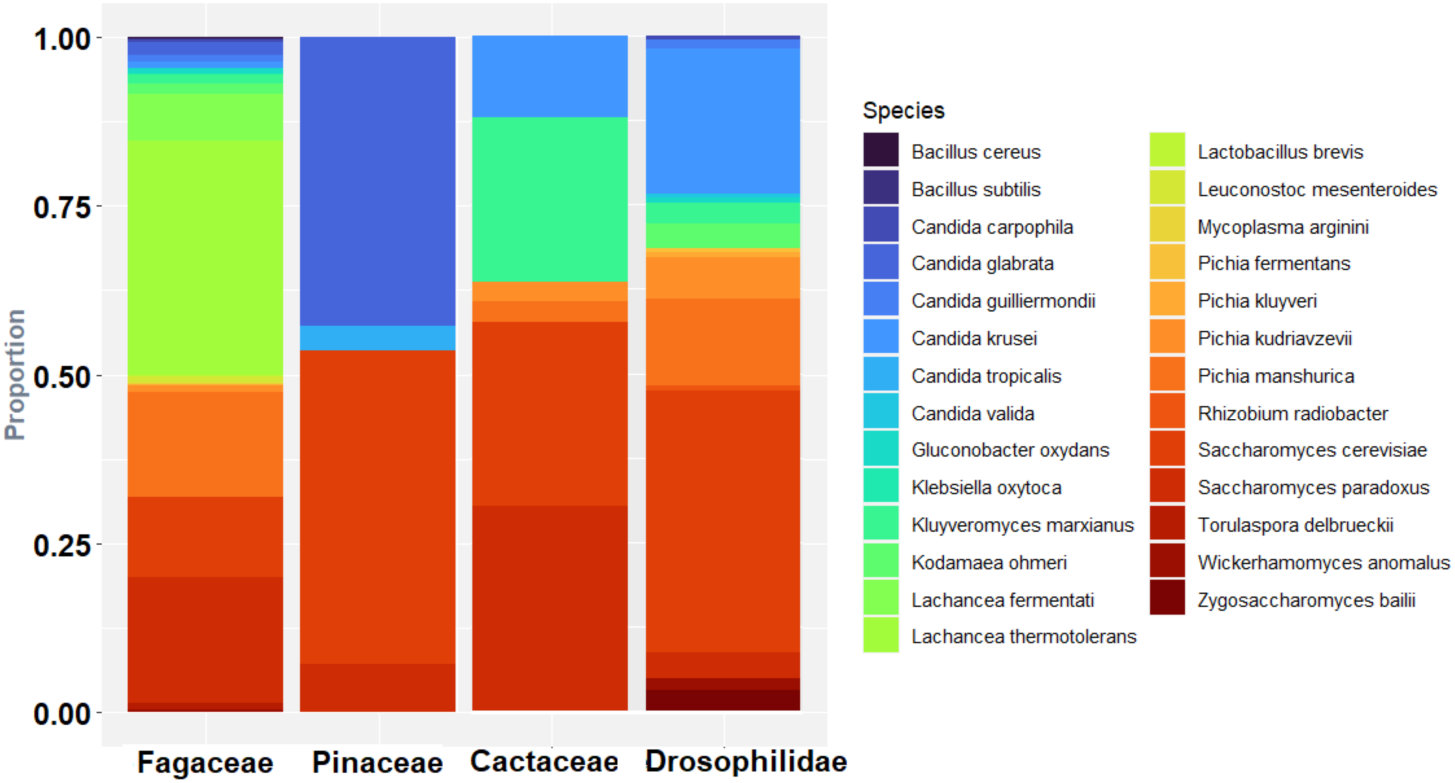
Diversity of species confidently identified with MALDI-TOF (score>1.5) isolated in plant families with the highest rate of isolation (Fagaceae, *n* = 166 samples and *n* = 494 isolates), Pinaceae (*n* = 46 samples, *n* = 28 isolates), Cactaceae (*n* = 5 samples, *n* = 33 isolates), and in the insect family Drosophilidae (*n* = 84 samples, *n* = 162 isolates).

**Suppl. Mat. S2.C.**
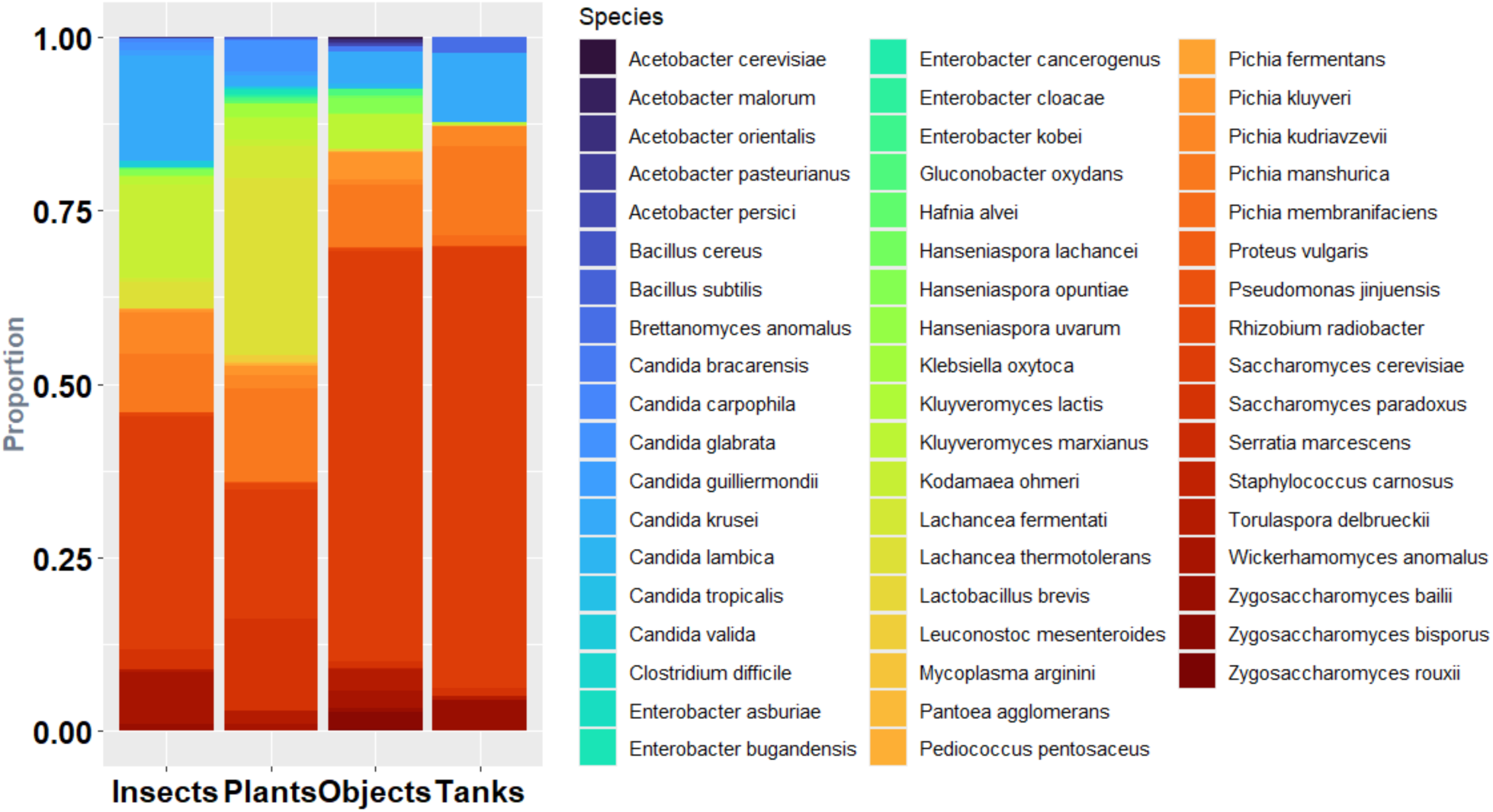
Diversity of species confidently identified with MALDI-TOF (score>1.5) isolated from different substrates regardless of location type. Insects (*n* = 341 samples, *n* = 517 isolates), plants (*n* = 382 samples, n=848 isolates), objects (*n* = 98 samples, *n* = 495 isolates), tanks (*n* = 40 samples, 349 isolates).

## SUPPLEMENTARY MATERIAL S3

**ADMIXTURE analyses of *Saccharomyces cerevisiae* strains from agave-spirits distilleries and natural sites of Mexico**

**Suppl. Mat. S3.A.**
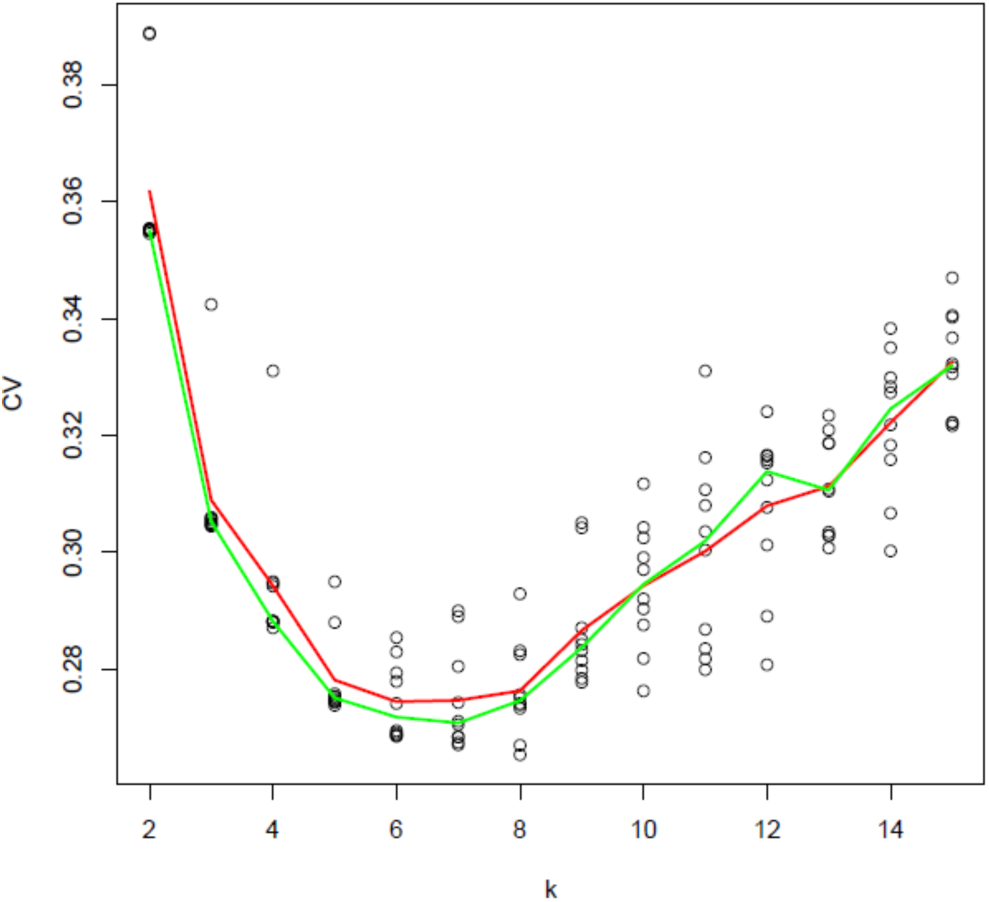
ADMIXTURE cross-validation errors for K=2 to K=15, in a matrix of 108,636 SNPs (previously LD pruned) and 88 *S. cerevisiae* strains. The red line represents the mean, whereas the green line represents the median.

**Suppl. Mat S3.B.**
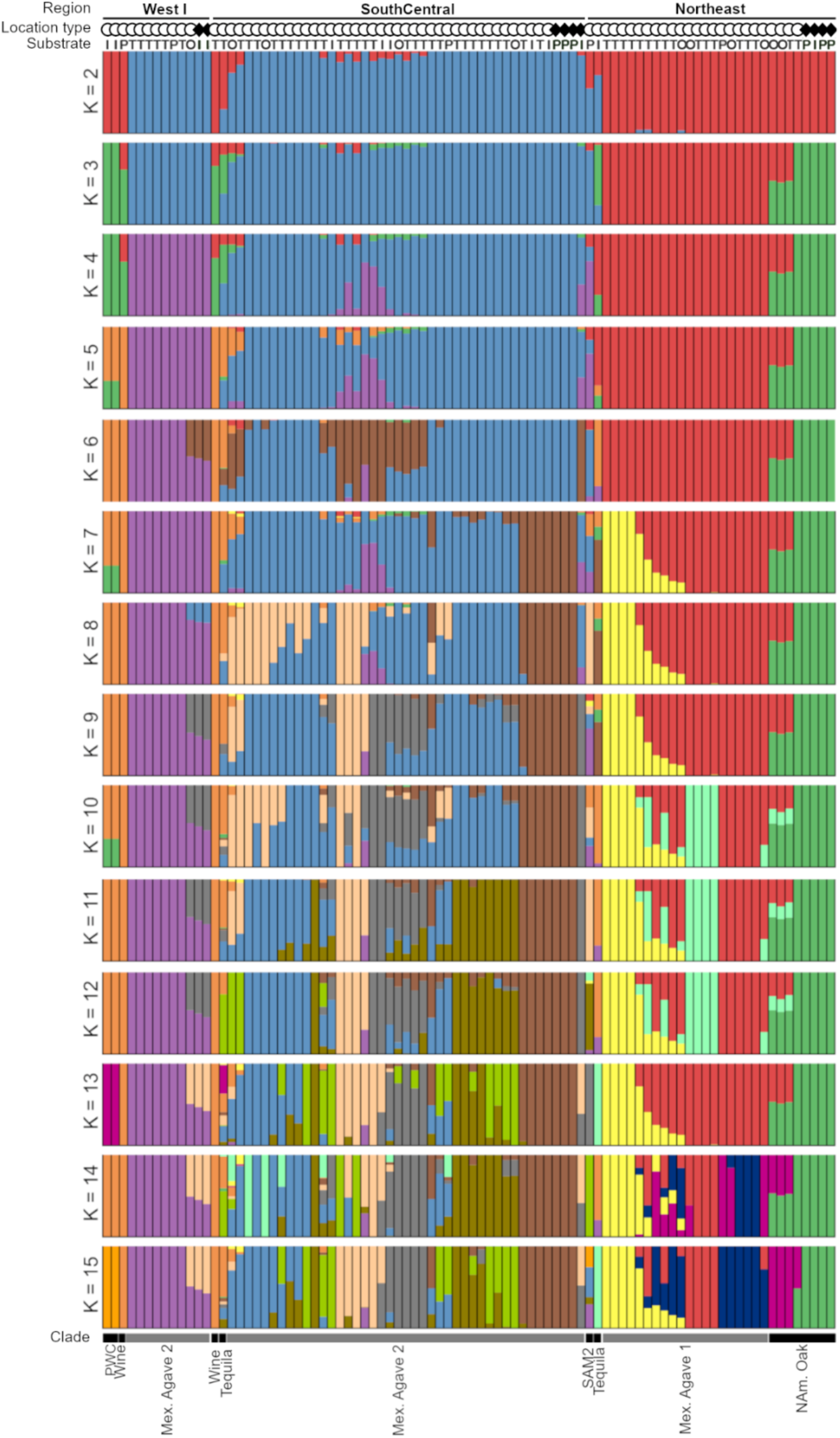
ADMIXTURE plots of 88 *S. cerevisiae* strains from *K* = 2 to *K* = 15.

## SUPPLEMENTARY MATERIAL S4

**ADMIXTURE analyses of *Saccharomyces paradoxus* strains from agave-spirits distilleries and natural sites of Mexico**

**Suppl. Mat. S4.A.**
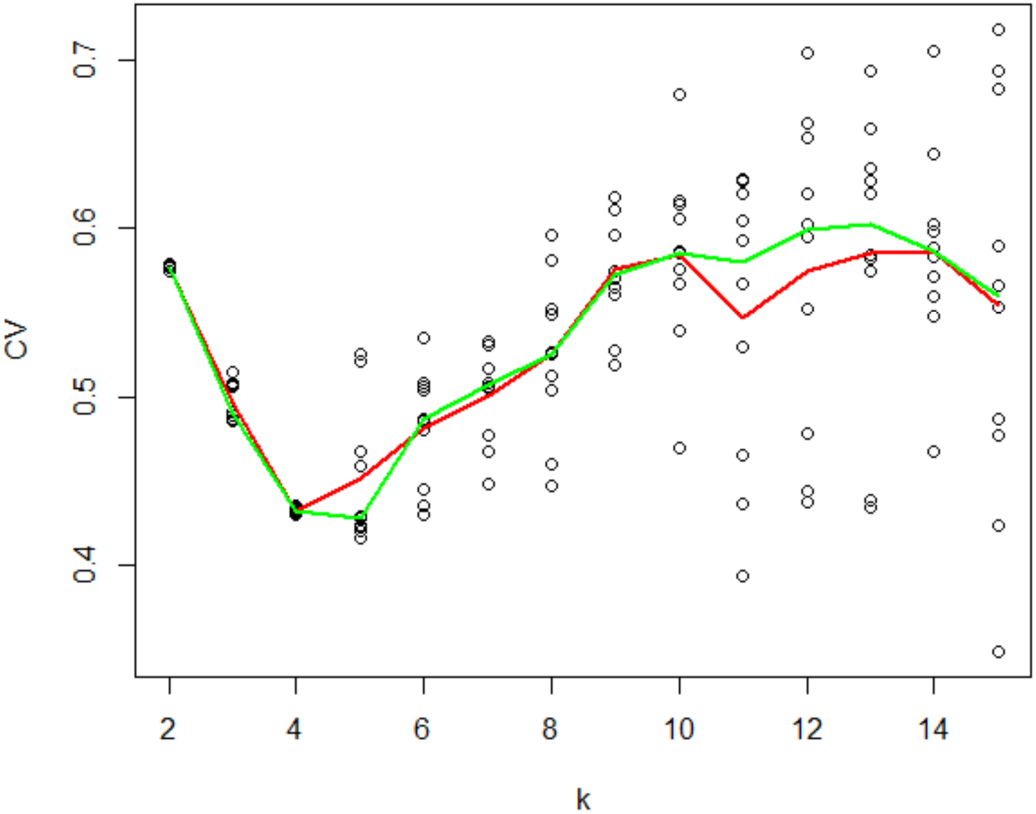
ADMIXTURE cross-validation errors for *K* = 2 to *K* = 15, in a matrix of 86,448 SNPs (previously LD pruned) and 47 *S. paradoxus* strains. The red line represents the mean, whereas the green line represents the median.

**Suppl. Mat. S4.B.**
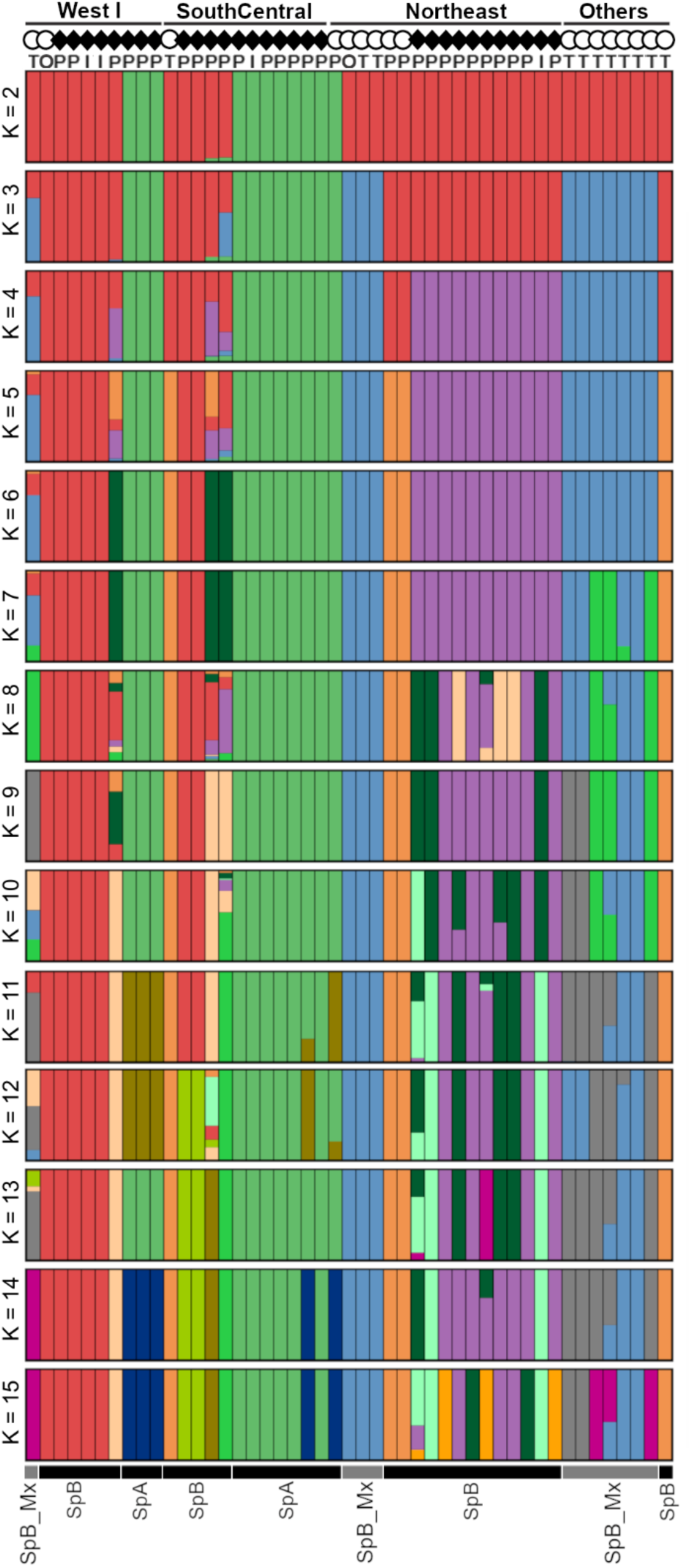
ADMIXTURE plots for 47 *S. paradoxus* strains from *K* = 2 to *K* = 15, with 86,448 SNPs (previously LD pruned).

## SUPPLEMENTARY MATERIAL S5

**Suppl. Mat. S5.A.**
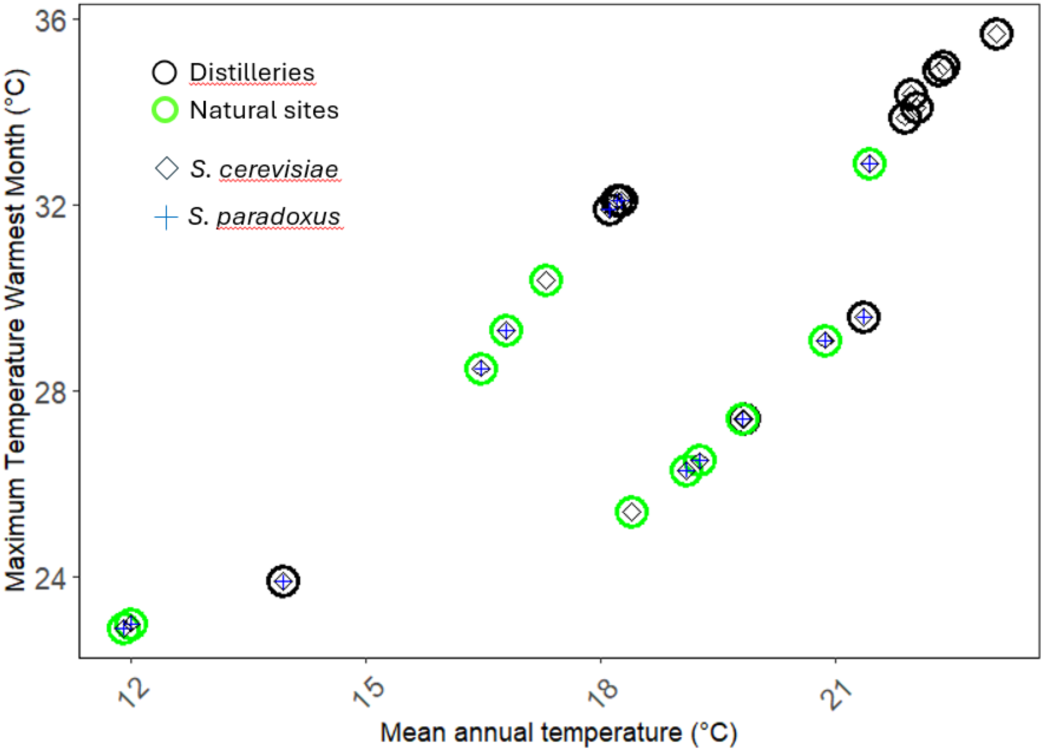
Presence of *Saccharomyces* species in distilleries and natural sites according to climatic conditions of sampled localities. Mean annual temperature and Maximum Temperature of Warmest Month at each location are based on WorldClim data (see Suppl. Mat. S1; Fick & Hijmans, 2017).

**Suppl. Mat. S5.B.**
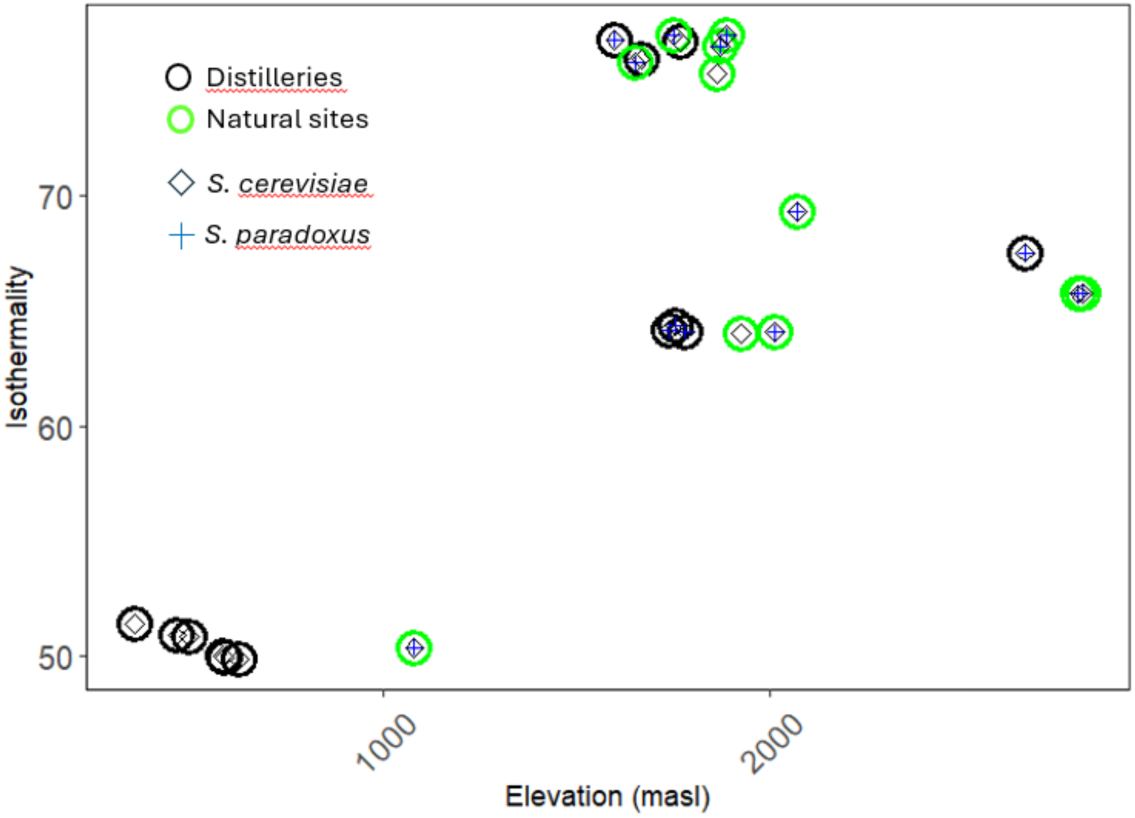
Presence of *Saccharomyces* species in distilleries and natural sites according to climatic conditions of sampled localities. Isothermality at each location is based on WorldClim data. Isothermality is a way of measuring how consistent the temperature is throughout the year, with higher values indicating daily temperature variations that remain relatively small when compared to temperature changes through seasons of the year (Fick & Hijmans, 2017).

